# Exploring evolutionary relationships across the genome using topology weighting

**DOI:** 10.1101/069112

**Authors:** Simon H. Martin, Steven M. Van Belleghem

## Abstract

We introduce the concept of topology weighting, a method for quantifying relationships between taxa that are not necessarily monophyletic, and visualising how these relationships change across the genome. A given set of taxa can be related in a limited number of ways, but if each taxon is represented by multiple sequences, the number of possible topologies becomes very large. Topology weighting reduces this complexity by quantifying the contribution of each ‘taxon topology’ to the full tree. We describe our method for topology weighting by iterative sampling of sub-trees (*Twisst*), and test it on both simulated and real genomic data. Overall, we show that this is an informative and versatile approach, suitable for exploring relationships in almost any genomic dataset.

Scripts to implement the method described are available at github.com/simonhmartin/twisst.

## Introduction

The relationship (or genealogy) among recombining DNA sequences from closely-related taxa often varies across the genome, due to both variation in lineage sorting and introgression (Maddison 1997). Numerous methods focus on inferring, from this genealogical variation, the underlying population branching pattern (the 'species tree') (e.g. Heled and Drummond 2010) or demographic history (e.g. Lohse et al. 2012). It is now also possible to characterise the complete genomic landscape of relatedness using whole genome sequences. For example, Hobolth et al. (2007) and Dutheil et al. (2009) developed an approach that uses whole genome sequences to infer not only the population history, but also how and where in the genome the genealogy changes. More recent studies have attempted to characterise patterns of relatedness along larger numbers of whole genomes, either by simply inferring phylogenetic trees for pre-defined windows (Martin *et al*. 2013; Fontaine *et al*. 2015), or by attempting to infer both the trees and the likely breakpoints that separate them (Gante *et al*. 2016). An emerging challenge with increasing numbers of sequences, is that both the inference and interpretation of genealogies becomes difficult, due to the rapid escalation of topological complexity. For example, for five haploid sequences, there are fifteen possible unrooted, bifurcating tree topologies, whereas for ten sequences there are over two million. Here we address this challenge of characterising and summarising the genomic landscape of relatedness in large datasets with multiple genomes from multiple taxa.

One way to deal with the problem of increasing tree complexity is to focus specifically on the relationships among broader predefined taxa (hereafter “taxon topologies”), and not among all of the sequences. This is straightforward if each taxon is completely resolved into a monophyletic clade, so that the branching patterns within each taxon can simply be ignored, but it becomes challenging when the taxa are not reciprocally monophyletic (i.e. when lineages are not completely sorted). This is often the case for closely-related taxa, in which lineages from the same population coalesce (share a most recent common ancestor) in the ancestral population, and may therefore be more closely related to lineages from other taxa than from their own (Fig. 1A). We note that tree inference using large genomic windows, entire chromosomes or whole genomes often yields completely-sorted monophyletic taxa, but this may be artificial as such methods are usually forced to infer a single best-supported tree, even though the region may represent multiple distinct incompletely-sorted ancestries. Population genetic statistics based on allele frequencies, such as *F*_ST_, have the advantage of *quantifying* relatedness rather than simply qualitatively describing the topology. However, by ignoring topology entirely, they become difficult to interpret when the number of taxa is greater than two. There is therefore a need for descriptive methods that incorporate both the qualitative tree-like structure of relationships and the quantitative variation in taxon relationships across the genome. Examples of such methods include the so-called ABBA-BABA test (Green *et al*. 2010; Durand *et al*. 2011; Martin *et al*. 2015), and *f* statistics (Reich *et al*. 2009, 2012; Patterson *et al*. 2012), both of which evaluate the support for alternative taxon topologies using allele frequencies at single nucleotide polymorphisms (SNPs). However, because individual SNPs are only informative about two separate groupings, these methods do not scale to more than four taxa.

**Figure 1.**
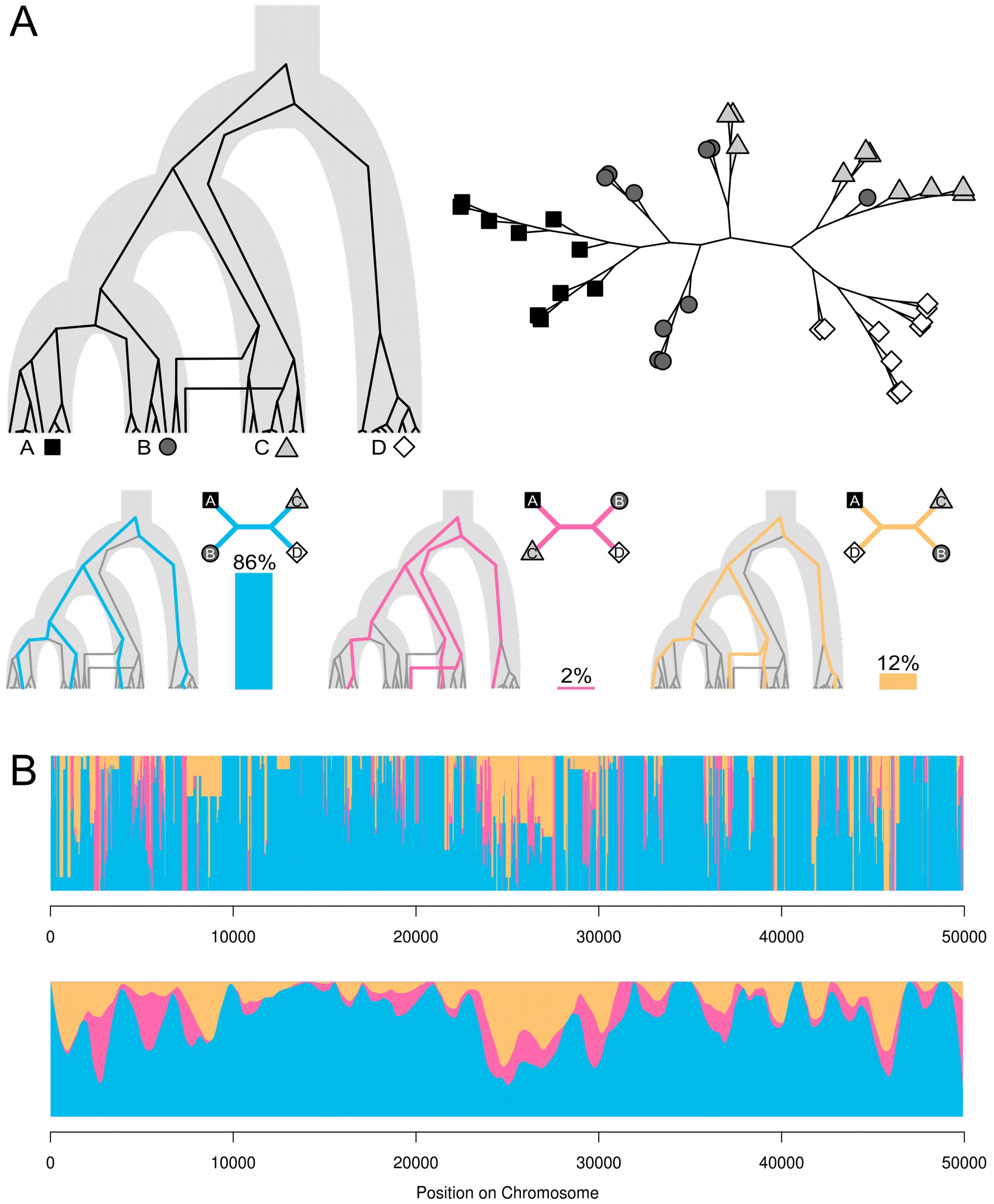
Topology weighting.

**(A)** An example genealogy for four taxa (A, B, C and D), plotted as an unrooted tree on the top right. Taxa B and C are not monophyletic, due to both deep coalescences causing incomplete lineage sorting and gene flow. The three possible taxon topologies are shown, along with a single example sub-tree that matches each topology. The percentage of all sub-trees matching each taxon topology (i.e. the weightings) are shown by vertical bars. **(B)** Topology weightings plotted across a 50 kb region of a simulated recombining chromosome. Weightings for the three topologies are stacked (they always sum to 1, as they are proportions). Changes in the weightings along the chromosome indicate regions of distinct genealogical history separated by recombinations. Below, the same data are plotted with loess smoothing (span = 2.5 kb).

Here we introduce the concept of topology weighting, which offers a simple and general solution to the problem of quantifying relationships among taxa that are not necessarily monophyletic. Given a tree of relationships for a set of taxa, each represented by an arbitrary number of sequences, topology weighting quantifies the contribution of each individual 'taxon topology' to the full tree. We describe our approach to compute topology weightings, which we call *Twisst* (topology weighting by iterative sampling of sub-trees), and explore the utility of this approach using simulated data as well as two different genomic datasets from butterflies and fungi. Overall, we show that this concept provides a useful means to explore relationships using genomic data, both to test hypotheses and generate new ones.

## Materials and Methods

### Topology weighting by iterative sampling of sub-trees

A given set of taxa can be related in a limited number of ways. For example, for four taxa labelled A, B, C and D, there are three possible unrooted bifurcating topologies: ((A,B),C,D), ((A,C),B,D) and ((A,D),B,C) (Fig. 1A). Given a tree with any number of tips (or leaves), each assigned to a particular taxon, we define the weighting of a particular 'taxon topology', *τ*, as the fraction of all unique sub-trees – in which each taxon is represented by a single tip – that match the particular taxon topology:

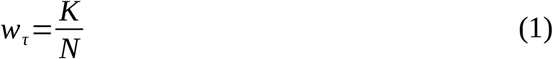

Where *N* is the number of possible unique sample sets in which each taxon is represented by a single sample. This corresponds to the product of the number of samples in each taxon. *K* is the number of these unique sample sets for which the corresponding sub-tree topology matches *τ*. Thus:

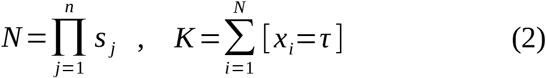

In which *n* is the number of defined taxa (groups), *s_j_* is the number of samples in taxon *j*, and *x_i_* is the sub-tree topology corresponding to subset *i*.

Topology weighting therefore reduces the complexity of the full tree to a number of values, each giving the proportionate contribution of a particular taxon topology (Fig. 1A). This method has conceptual similarities to the quartet sampling approach for comparing topologies (Estabrook *et al*. 1985), except that here we do not consider all sub-trees, but only those in which each tip represents a different taxon. Moreover, topology weighting can be applied to any number of taxa. The taxa can be defined arbitrarily, for example by phenotype or geography, as with the 'operational taxonomic units' used in biogeography. Because the number of taxon topologies is limited, the weightings can be normalised to sum to 1, making them easily comparable between different parts of the genome.

We computed topology weightings using our *Twisst* approach, implemented by Python scripts available for download at (https://github.com/simonhmartin/twisst). The *Twisst* algorithm first computes all possible unrooted, bifurcating taxon topologies, and then determines the number of unique sub-trees that match each topology by iteratively sampling a single individual from each taxon and 'pruning' away all other branches and nodes. Our implementation makes use of the Environment for Tree Exploration, ETE v3 (Huerta-Cepas *et al*. 2016).

The number of unique sample combinations (and corresponding sub-trees) can be very large. However, the iterative process is sped up considerably by first collapsing monophyletic groups of samples from the same taxon and weighting these nodes proportionately (Fig. S1). Nevertheless, if the taxa are highly unsorted and the tree is large, it may not be possible to consider all possible sample combinations in a reasonable amount of time. In such cases, approximate weightings can be computed by randomly sampling a subset of sample combinations (Fig. S2). Random sampling for approximate weighting is performed with replacement, so that the errors fit a binomial distribution, allowing for the computation of a confidence interval (Fig. S2). Our implementation of *Twisst* also allows a threshold-based sampling procedure, in which sampling is repeated until a particular level of confidence is achieved. This allows further speed-ups since highly sorted trees, in which weightings are biased toward one or a few topologies, require less sampling for high confidence than unsorted or 'star-like' trees where most weightings are intermediate. For all analyses in this study, dataset sizes allowed for computation of complete weightings.

### Analysis of simulated chromosomes

In order to test our method on realistic data for which the complete genealogical history is known, we simulated the evolution of recombining chromosomes using the coalescent simulator *msms* (Ewing and Hermisson 2010). Simulations involved four taxa, each represented by 10 haploid sequences (40 in total), that split in the order (((A,B),C),D) (Fig. 2A), or five taxa, each represented by 6 sequences, that split in the order (((A,B),(C,D)),E) (Fig. S7A), with split times of 0.5, 1 and 1.5 (in units of 4*N* generations). Population size was constant throughout. In all scenarios, unidirectional migration from C to B was simulated. The simulation was performed for a 1 Mb chromosome, with a population recombination rate (4*Nr*) of 0.01 or 0.001. Genealogies for each unique ancestry block (separated by recombinations) were recorded. These were used to calculate the 'true' weightings using *Twisst*.

**Figure 2.**
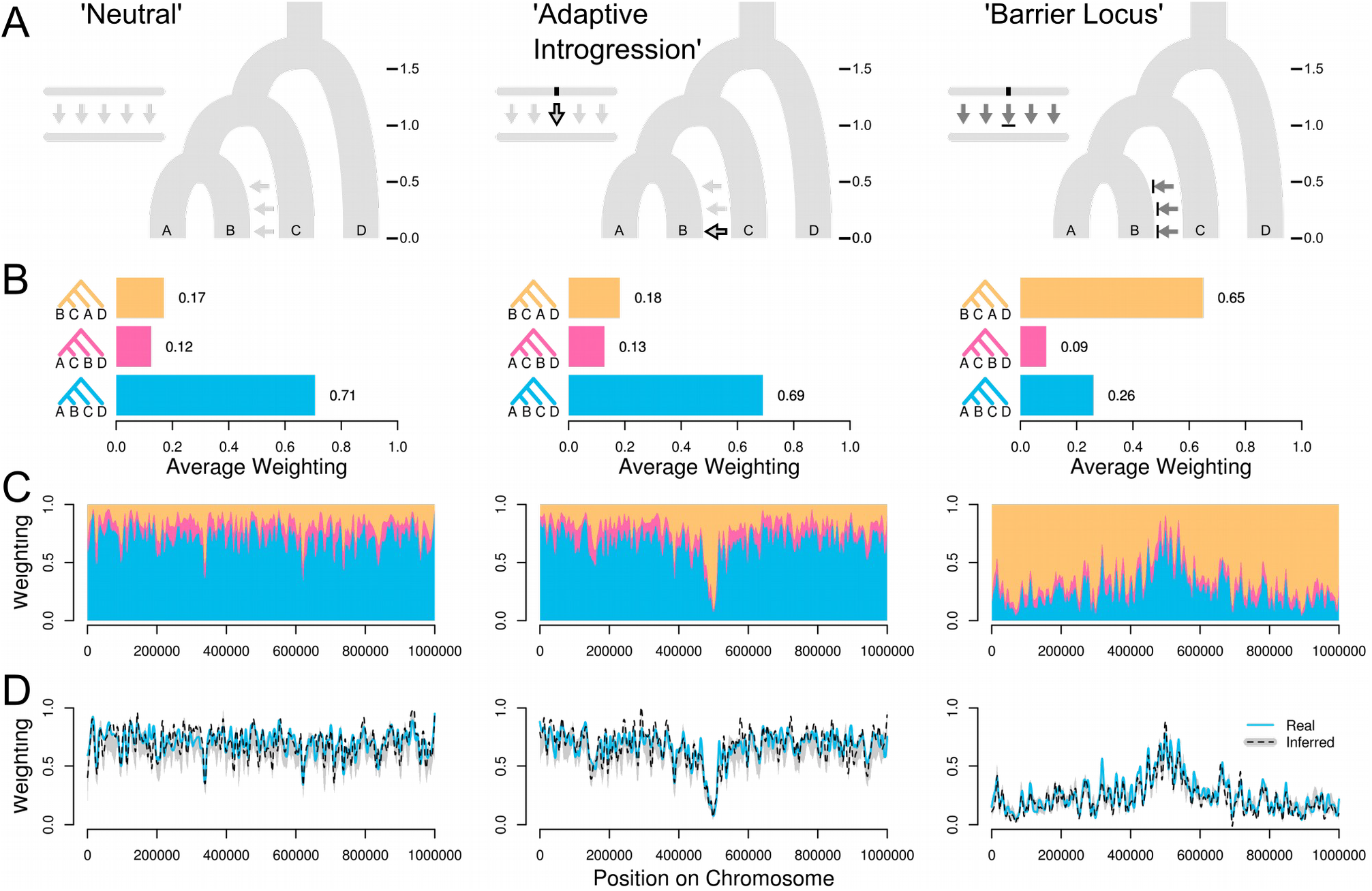
Tests on simulated chromosomes. **(A)** In all three demographic scenarios, populations split in the order (((A,B),C),D), at the split times indicated (in units of 4N generations), with migration from C to B. In the ‘Neutral’ scenario, there is no selection and moderate migration. The ‘Adaptive Introgression’ scenario is similar, except a beneficial allele at a locus in the centre of the chromosome is allowed to move from population C into B at time 0.1. In the ‘Barrier Locus’ scenario, the rate of migration is high, but an allele at the central locus that is fixed in C is selected against in population B. **(B)** Mean weightings for the three possible taxon topologies across the 1 Mb simulated chromosome. Note that we illustrate the topologies for the four taxa as rooted, with D as the outgroup for simplicity, but the rooting is not considered when computing the weightings. **(C)** Weightings for all three topologies plotted (stacked) across the chromosome, with loess smoothing (span = 20 kb). **(D)** Weightings for topology (((A,B),C),D) inferred from simulated sequence data using non-overlapping 50 SNP windows and neighbour joining. Solid blue lines indicate the true values, and dashed black lines indicate the inferred values. Grey shading indicates the lower (5%) and upper (95%) quantiles based on 100 bootstrap replicates. Values are smoothed as above.

Three distinct evolutionary scenarios were simulated (Fig. 2A, Fig. S7A). *msms* command options are provided in Supplementary Text 1. The first was a 'Neutral' scenario, with no selection and a low migration rate from B to C of 0.1 (in units of 4*Nm*, where *m* is the fraction of B made up of migrants from C each generation). The second was an 'Adaptive Introgression' scenario, which is the same as above except that a beneficial allele at a locus in the centre of the 1 Mb chromosome is allowed to move from population C into B at time 0.1. This was achieved by initiating selection at this time point on a dominant allele that was fixed in population C and absent from the other populations. A selection coefficient of 0.005 was used for both the homozygote and heterozygote, with a diploid population size of 100,000, giving a selection strength (2*Ns*) of 1000 for both genotypes. The third was a 'Barrier Locus' scenario, where the rate of migration from C to B was 5 (in units of 4*Nm*, as above), and a dominant allele at the central locus that is fixed in C is selected against in population B. The same selection coefficient and population size as above were used.

We simulated sequences from the simulated genealogies using *seq-gen* (Rambaut and Grass 1997). Command options are provided in Supplementary Text 1. The branch scaling factor for mutation was 0.01. Since branches in the simulated genealogies are in units of 4*N* generations, taking *N* to be 100,000 gives μ (the per generation mutation rate) of 2.5 × 10^−8^.

### Inferring trees in sliding windows

We tested different methods for inferring trees in windows across the chromosome. The simplest approach used non-overlapping windows of a fixed number of SNPs. A range of window sizes were tested. Trees were then inferred for each window using PhyML version 3.0 (Guindon *et al*. 2010), implementing either the BIONJ neighbour-joining algorithm (Gascuel 1997) or maximum likelihood optimization of the topology and branch lengths. To investigate the consistency of the tree inference and compute confidence thresholds for the weightings, we performed bootstraping by randomly re-sampling the SNPs in each window with replacement and repeating the tree inference. We also tested an approach to infer likely window breakpoints from the data. Taking the topology weightings computed from 10 SNP windows, we used the R package *GenWin* (Beissinger *et al*. 2015) to fit a beta-spline to the data and find likely inflection points, which we then used as window breakpoints and inferred a new set of trees for these. In addition, we tested the program *Saguaro*, which infers both the breakpoints and the distance matrix describing each region. Distance matrices were converted to trees using BIONJ, as above.

### Power Analyses

An important aspect of our approach is its dependence upon reliable trees, which may be inferred from relatively short sequence windows. To investigate the power we have to infer topology weightings from short sequences, we simulated datasets under a range of sampling strategies and demographic scenarios, and then compared the true weightings to those computed using trees inferred from the simulated sequences.

Eight sampling strategies were compared, including four, five, six or ten sequences from either four or five populations (Fig. S3). For each sampling strategy, two different demographic scenarios were simulated. In the four-population scenarios, the populations split in the order ((1,2),(3,4)), with the basal split time (*t1*) at either 0.5 or 1 × 4N generations in the past, and the splits between populations 1 and 2 and 3 and 4 both occurring at 0.1 × 4N generations in the past (*t2*) (Fig. S3). In the five-population cases, the populations split in the order (((1,2),(3,4)),5). As above, the basal split time (*t1*) occurs at either 0.5 or 1 × 4N generations in the past. The next split, between populations 1 and 2 and populations 3 and 4, occurs at 0.2 × 4N generations in the past (*t1b*), and the final two splits between populations 1 and 2, and 3 and 4 both occur at 0.1 × 4N generations in the past (*t2*) (Fig. S3).

In each run, we used *msms* (Ewing and Hermisson 2010) to simulate 500 genealogies for the given sampling design and demographic scenario. For each genealogy, we computed the topology weightings using *Twisst*, and then generated a simulated set of sequences using *seq-gen* (Rambaut and Grass 1997). The sequences were then truncated at different lengths to compare tree inference using 10, 25, 50, 100, 200 or 400 SNPs. Trees were inferred in Phyml (Guindon *et al*. 2010), using either BIONJ, or with maximum likelihood optimisation of the topology and branch lengths. Weightings were then computed from the inferred tree, and compared to the set of true weightings using a scaled euclidean distance:

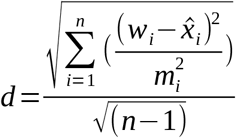

Where *n* is the number of weighting values (i.e. the number of taxon topologies), *w_i_* is the true weighting for topology *i* and *x̂_i_* is the inferred weighting for topology *i*. *m_i_* is the absolute value of the maximum possible distance from *w_i_*. This value therefore gives a distance between the true and inferred sets of weightings on a scale of 0 to 1, with 0 indicating identical values (i.e. perfect inference) and 1 indicating a maximum possible discrepancy between the true and inferred values.

Not all SNPs are phylogenetically informative, and even those that are (those that are not singletons), are not necessarily informative about the relationships among the broader taxa, which is of primary interest for topology weighting. We therefore also tested the power of inference using a subclass of Taxon Informative Sites' (TISs), which we define as having at least two alleles present in at least two taxa. As above, simulated sequences were truncated to contain the number of TISs.

### Analysis of real genomic data

We tested *Twisst* on two published genomic datasets from *Neurospora* spp. (ascomycete fungi) and *Heliconius* spp. (butterflies), selected to represent different sampling strategies (4 and 5 taxa, respectively), as well as different levels of evolutionary complexity. The *Neurospora* dataset (Corcoran *et al*. 2016) consisted of 22 aligned haploid genome sequences from *Neurospora tetrasperma* samples (10 of mating type *A*, and 12 of mating type *a*), along with single genomes representing two related species: *Neurospora crassa*, and *Neurospora hispaniola*. Whole genome alignments were obtained from datadryad.org/resource/doi:10.5061/dryad.162mh. We used Lineage-10 (UK) samples of *N. tetrasperma*, as these had been shown to carry a strong signal of introgression from *N. hispaniola* (Corcoran *et al*. 2016). Trees were constructed for sliding windows of 50 SNPs using BIONJ as described above, with the requirement that each sample had to be genotyped at 40 or more of the 50 SNPs per window. Topology weightings were computed using *Twisst*, with four defined taxa: *N. tetrasperma mat a* (12 sequences), *N. tetrasperma mat A* (10), *N. crassa* (1) and *N. hispaniola* (1).

The *Heliconius* dataset consisted of 18 resequenced genomes (or 36 haploid genomes) from Martin et al. (Martin *et al*. 2013). These samples comprised five populations: two geographically isolated races of *Heliconius melpomene*, from Panama (*H. m. rosina*, n=4) and Peru (*H. m. amaryllis*, n=4), and their respective sympatric relatives *Heliconius cydno chioneus* from Panama (n=4) and *Heliconius timareta thelxinoe* from Peru (n=4), with which they are known to hybridize; along with two additional samples of the more distant 'silvanifrom' clade to serve as outgroups. We limited our analysis to two chromosomes: 18, which carries the gene o*ptix*, known to be associated with red wing pattern variation; and 21, the Z sex chromosome, which has been shown to experience reduced gene flow between these species, probably due to genetic incompatibilities (Martin *et al*. 2013). Fastq reads were downloaded from the European Nucleotide Archive, study accession ERP002440. Reads were mapped to the *Heliconius melpomene* reference genome v2 (Davey *et al*. 2016) using BWA-mem (Li and Durbin 2009; Li 2013) with default parameters. Genotyping was performed using the Genome Analysis Toolkit (DePristo *et al*. 2011) v3 HaplotypeCaller and GenotypeGVCFs, with default parameters except that heterozygosity was set to 0.02. Phasing and imputation was performed using Beagle v4 (Browning and Browning 2007). Topology weighting was performed using *Twisst* using the five taxa described above.

## Results

### Analysis of simulated chromosomes

Topology weighting provides an informative summary of the genealogical data, and highlights differences between the simulated scenarios (Fig. 2). As described above, there are three possible unrooted topologies for the four taxa. In the 'Neutral' scenario, the most prevalent topology, (((A,B),C),D), which reflects the population split times, has an average weighting of 71% across the chromosome. The other two topologies are both fairly rare, but one (((B,C),A),D) is more common on average (17%) than the other (((A,C),B),D) (12%). This is because the former can result from both gene flow and incomplete lineage sorting (ILS), whereas the latter can only result from ILS, as there was no simulated migration between A and C or between B and D. In the 'Adaptive Introgression' scenario, the weightings are very similar to the Neutral scenario on average, but in the centre of the chromosome there is a strong excess of the topology (((B,C),A),D), created by the spread of a beneficial allele from population C into B. Finally, in the 'Barrier Locus' scenario, high migration from C to B causes a swamping by the topology (((B,C),A),D), which has an average weighting of 65%. However, there is a broad peak at the centre of the chromosome where the population branching topology (((A,B),C),D) had not been eroded, due to selection limiting introgression.

In the corresponding simulations with five taxa, there are fifteen possible taxon topologies (Fig. S7). There is greater topological variation overall, as there are more ways that incomplete sorting can occur. Nonetheless, topology weights clearly detect the differences among the scenarios, highlighting the most abundant topologies as well as the location of the selected locus (Fig. S7).

### Inferring weightings from simulated sequence data

Above, we computed the weightings directly from the simulated genealogies, but we are also able to show that topology weightings can be reliably estimated when the genealogies are inferred from simulated sequence data (Fig. 2D, S7D). Because neither the genealogies, nor the recombination breakpoints at which genealogies switch are known, we tested several approaches for inferring genealogies for narrow intervals across the chromosome. First, we performed extensive power analyses, covering a range of demographic scenarios and sampling designs, to explore the relationship between the number of SNPs used for tree inference and the accuracy of topology weighting. Across the range of scenarios investigated, we find a consistent lower bound of 50 SNPs to achieve >90% accuracy (Fig. S4, S5, S6). Focussing specifically on 'taxon informative sites' (see above) makes no discernible difference, probably because most SNPs in our simulations are taxon informative. These tests also indicate that neighbour-joining trees provide more accurate weightings than maximum likelihood trees, in addition to much faster computation (Fig. S4, S5, S6).

We then analysed trees inferred for non-overlapping windows across our simulated recombining chromosomes. A fixed window size of 50 SNPs gives results that most closely approximate the true weightings (Fig. 2D, S7D). In agreement with our power analyses, with fewer than 50 SNPs, the estimates are less accurate, and tend to underestimate the weighting of the most prevalent topology (Fig. S8, S9). Weightings tending toward intermediate values are expected as the underlying trees become less well resolved. Interestingly, windows of 100 SNPs or above also result in reduced accuracy, but with a tendency to overestimate support for the most prevalent topology and underestimate support for others (Fig. S8, S9). This can be explained by the fact that large windows are forced to average over regions of distinct ancestry, therefore favouring the most widespread signal. To confirm this hypothesis, we repeated our neutral simulation using a ten-fold lower population recombination rate. In this new dataset, 100 SNP windows give the most accurate weightings, and even 200 SNP windows have high accuracy, while 50 SNP windows perform only marginally less well (Fig. S10, S11).

We tested whether bootstrapping over the SNPs in each window can be used to validate the accuracy of the observed weightings. Bootstrap weights tend to be similar, but marginally more conservative, underestimating the weight of the most prevalent topology (Fig. 2D). This is because the bootstrap trees tend to be slightly less well resolved, leading to more intermediate weightings. Bootstrapping is therefore a useful means to test the strength of support for an observed peak in the weighting of a particular topology. However, being inherently conservative, bootstrapping would not be able to determine whether an observed intermediate weighting was accurate or simply the result of a poorly resolved tree.

Because real recombination breakpoints are not evenly spaced, we also tested two approaches in which the window boundaries are inferred from the data itself. In our first approach, we used the R package *GenWin* (Beissinger *et al*. 2015) to fit a smooth spline to the weightings from 10-SNP windows and identify likely window boundaries as inflection points, and then inferred trees for the the newly-defined window regions. The resulting topology weightings match the true weightings fairly well, but not as well as for the fixed 50 SNP windows (Fig. S12, S13). As above, this appears to be due to poor tree inference in the smallest windows. The second approach used the method *Saguaro* (Zamani *et al*. 2013), which combines a Hidden Markov Model and a Self Organising Map to infer both the trees and window boundaries. This approach poorly recapitulates the true weightings, greatly overestimating support for the most prevalent topology (Fig. S12, S13). We therefore used fixed windows of 50 SNPs for all further analyses.

### Branch lengths differ among topology types

Topology weighting is primarily a descriptive method, but the weightings do carry information that can aid inferences about population history. The simulated ‘Barrier Locus’ scenario (Fig. 2) provides an interesting test case. Due to the overwhelming signal of introgression, it would be difficult to know which topology corresponds to the true population branching order (i.e. the ‘species tree’) if this was not known. The topology (((B,C),A),D) is prevalent across much of the chromosome, but (((A,B),C),D) is prevalent in around the chromosome centre. It has been proposed that the original population branching order can be identified by considering branch lengths (Fontaine *et al*. 2015; Gante *et al*. 2016). Taxa that cluster together due to recent introgression tend to be separated by short branches, whereas those that cluster according to the population branching order should have deeper splits. Indeed, in trees inferred from 50 SNP windows, pairwise branch distances between the taxa suggest that sub-trees matching (((B,C),A),D) tend to result from recent introgression between B and C (Fig. S14), thus implying that (((A,B),C),D) is the more likely population branching order.

### Analysis of real genomic data

The *Neurospora* dataset consists of four taxa (three possible topologies), and is the simpler of the two real datasets analysed (Fig. 3A,B). It was selected to test how well *Twisst* is able to detect the signal of a previously described adaptive introgression event from *N. hispaniola* into *N. tetrasperma* individuals of the *A* mating type (Corcoran *et al*. 2016). This introgression covers the entire (∼7 Mb) non-recombining region of linkage group I (LGI). Indeed, we find a dramatic shift in the pattern of topology weightings in the central part of LGI (Fig 3C). The 'species-tree' topology (topo1), which groups the two *N. tetrasperma* mating types as closest relatives, is prevalent across most of the genome, but has very little weighting in the central part of LGI. Instead, it is replaced by topo3, which groups mating type *A* individuals of *N. tetrasperma* with *N. hispaniola*. Elsewhere, topo3 has limited weighting, nearly identical to that of topo2, consistent with a low level of incomplete lineage sorting throughout the genome. However, a region of LGIV also shows a weak shift in support towards topo3, potentially reflecting a separate introgression signal involving a small number of sequences.

**Figure 3.**
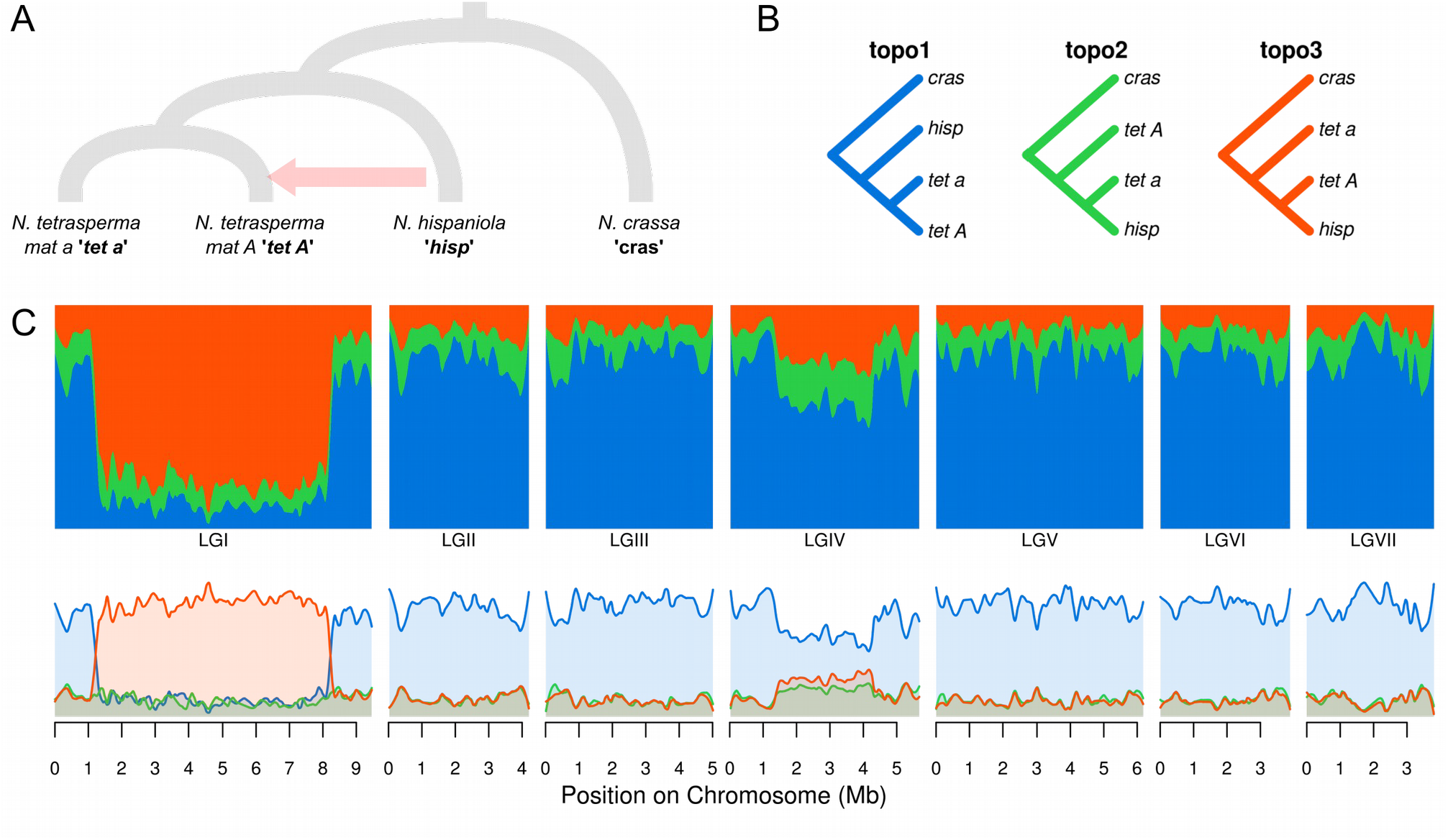
*Neurospora* analysis. **(A)** The putative species tree. Note that mating type *a* and *A* individuals of *N. tetrasperma* are shown as separate branches, while in reality, apart from the non-recombining region of LGI, these samples represent a single recombining population. The putative introgression from *N. hispaniola* into *N. tetrasperma mat A* individuals (Corcoran *et al.* 2016) is indicated. **(B)** The three possible taxon topologies for these four taxa. **(C)** Topology weightings for 50 SNP windows plotted across all seven linkage groups, with loess smoothing (span = 500 kb). The upper and lower plots show the same data, plotted as stacked or as separate lines, respectively.

The *Heliconius* dataset represents a more complex, five-taxon test case. The five taxa include an outgroup and two pairs of sympatric, non-sister taxa, between which gene flow is known to occur (Fig. 4A). Of the 15 possible topologies (Fig. 4B), the two most common across these chromosomes are topo3 and topo6. topo3 is consistent with the accepted species branching order, in which the allopatric *H. cydno chioneus* and *H. timareta thelxinoe* are sister taxa; whereas topo6 groups populations by geography, consistent with inter-specific gene flow in both Panama and Peru. The former is by far the most prevalent throughout the Z chromosome (Fig. 4C). By contrast, the species topology has variable weighting across chromosome 18, and is in places outweighed by topologies consistent with gene flow (topo4, topo5, topo6, topo11, topo14). In particular, there is a strong peak in the region of *optix* for topo11, which groups the taxa by wing pattern, and is consistent with the previously described adaptive introgression of the red-band allele between *H. m. amaryllis* and *H. t. thelxinoe* in Peru (Pardo-Diaz *et al*. 2012; The Heliconius Genome Consortium 2012). Zooming in on this peak shows a clear block of ∼150 kb over which the introgression topology is weighted highly (Fig. S16). This block includes the regulatory region downstream of *optix* that is known to controls wing pattern variation in these species (Baxter *et al*. 2010; Wallbank *et al*. 2016). Another four topologies that partially match the species branching order (topo1, topo2, topo10, topo15), have moderate weightings throughout, whereas topologies consistent with neither the species tree nor gene flow (topo7, topo8, topo9, topo12, topo13) have low weightings, especially across the Z chromosome, implying less incomplete lineage sorting than on chromosome 18.

**Figure 4.**
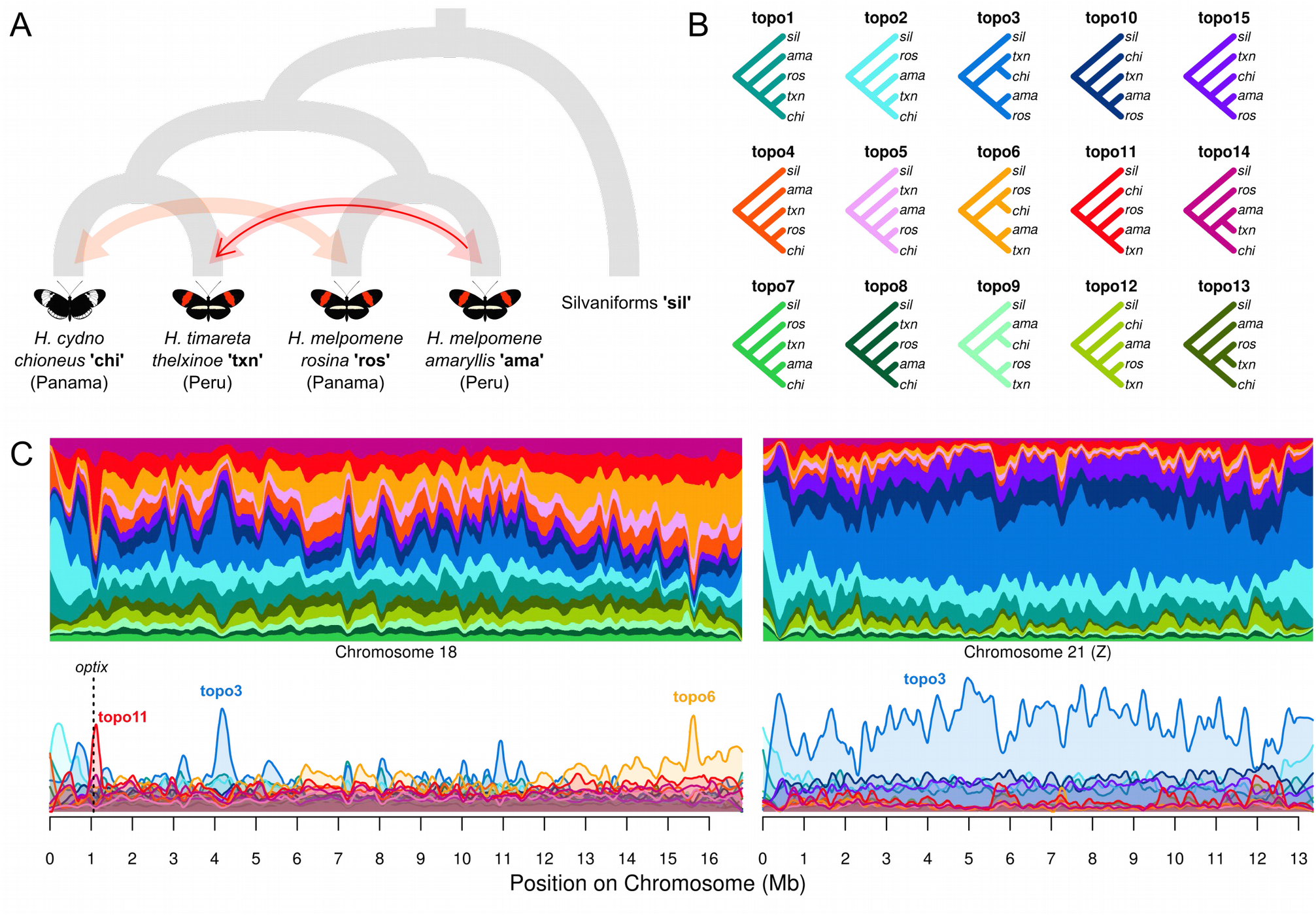
*Heliconius* analysis. **(A)** The putative species tree. Shaded arrows indicate ongoing gene flow between sympatric, non-sister taxa in Panama and Peru, respectively (Martin *et al.* 2013). The solid red arrow indicates the putative adaptive introgression of the the red wing-patterning allele near the gene *optix* (Pardo-Diaz *et al.* 2012; The Heliconius Genome Consortium 2012). **(B)** The fifteen possible taxon topologies for these five taxa. **(C)** Topology weightings for 50 SNP windows plotted across chromosomes 18 and 21 (Z), with loess smoothing (span = 500 kb). The upper and lower plots show the same data, plotted as stacked or as separate lines, respectively. The location of *optix* on chromosome 18 is indicated by a dashed vertical line.

## Discussion

Most statistics used in population genetics describe aspects of the underlying genealogy. For example, *F*_ST_ can be expressed as the relative rate of coalescence within sub-populations compared to the total population (Slatkin and Voelm 1991). Similarly, the *D* statistic of the ABBA BABA test compares the relative rate of coalescence between two pairs of non-sister populations (Green *et al*. 2010; Durand *et al*. 2011). Topology weighting can be seen as a generalization of this principle, as it determines the relative frequency of all possible patterns of coalescence among samples from a set of defined taxa. Unlike the ABBA BABA test, which is based on binary trees (samples either share the same allele or not) and therefore only four taxa, topology weighting uses a full genealogy, and can in principle be applied to any number of taxa, each represented by any number of sequences. In practice, however, beyond six taxa, the number of possible taxon topologies becomes very large, making topology weighting less practical. Nevertheless, even when the number of taxon topologies is very large, there may be value in comparing the weightings of particular topologies that support specific hypotheses (Van Belleghem *et al*. 2017). Unlike other methods for comparing tree topologies (e.g. Robinson and Foulds 1981; Estabrook et al. 1985), topology weighting reduces the problem to quantifying relationships among but not within defined taxa. There is also no real limitation on the number of samples included per taxon. Although computation of exact weightings may become infeasible for large trees, it remains possible to compute approximate weightings with small margins or error fairly rapidly. Our computational approach, *Twisst*, is based on a simple counting procedure, but we are confident that more efficient analytical solutions – or at least approximations – will be found.

An important consideration when applying this method is its dependence on the trees used. In most cases, the true genealogy for each distinct ancestry block is not known, and must be inferred from the sequences. Accurate inference requires multiple informative SNPs. Our tests on simulated data highlight a central difficulty when analysing recombining chromosomes: a trade-off between signal and resolution. Using larger numbers of SNPs increases our ability to infer the correct tree, but may average over genomic regions with different histories. This leads to a systematic overestimation of the weightings for more abundant topologies, whose signal tends to swamp out that of others. Using few SNPs per window allows for better resolution, but can lead to inaccuracies in tree reconstruction from insufficient signal (i.e. phylogenetic error), producing star-like trees and intermediate weightings for all topologies. Fortunately, this means that errors in tree inference are unlikely to result in spurious peaks in the weighting of a single topology, but instead may lead to underestimation of the height of a particular peak. A peak can be validated using bootstrapping, but an even better test is to demonstrate that it persists with decreasing window sizes. Our simulations, based on realistic recombination and mutation rates, indicate that a window size of 50 SNPs provides a good compromise between signal and resolution across a range of demographic scenarios, although larger windows may be acceptable if the population recombination rate is known to be low. While in some cases with high recombination rates, there may be too few mutations per recombination to accurately infer variation in genealogies across the genome, we expect that many cases will fall within a feasible range. Importantly, not all recombinations are relevant for topology weighting: only recombination events between lineages from distinct taxa (i.e. 'effective recombinations' or 'inter-taxon recombinations') can alter taxon relationships, and hence alter the weightings. The ability to infer the patterns of topology weighting across the genome therefore depends on the relationship between the mutation rate and the rate of inter-taxon recombination. Where possible, simulations tailored to the taxa being studied can be used to guide the choice of window size. In the future, improved methods to infer breakpoints from the data may further resolve this difficulty.

Another challenge is that diploid resequencing data should ideally be phased, so that each tip in the tree represents a distinct haplotype. Phasing can be performed using probabilistic approaches informed by patterns of linkage disequilibrium (Browning and Browning 2007; Delaneau *et al*. 2013). Although such methods may be error-prone across large genomic distances, they have fairly high accuracy at short ranges (Bukowicki *et al*. 2016), making them suited to the narrow windows used for topology weighting. Moreover, the genomic regions involving inter-taxon recombinations (i.e. those relevant for topology weighting) are more likely to be phased correctly, because they tend to involve more divergent sequences.

Topology weighting is principally a descriptive method, and can be applied with no prior knowledge of the studied samples, apart from some basis on which to define distinct groups, such as geography or phenotype. By capturing the tree-like nature of sequence evolution, it provides information that is not provided by descriptive statistics like *F*_ST_, or clustering methods such as *Structure* (Pritchard *et al*. 2000). In addition to describing the taxon branching order, tree-based methods allow incorporation of additional parameters like a nucleotide substitution model and different rates of evolution in different parts of the tree. Unlike conventional phylogenomic methods, topology weighting captures information about fine-scale and quantitative variation across the genome. This power and resolution is highlighted in the *Heliconius* example studied here, where topologies supporting admixture are common across chromosome 18, but there is one narrow peak consistent with the adaptive introgression of a wing patterning allele near the gene *optix*, as described previously (Pardo-Diaz *et al*. 2012; The Heliconius Genome Consortium 2012). We note that topology weighting simply describes the signal in the data, and does not explicitly test for introgression over other causes of discordant phylogenetic signal. For example, the elevated frequency of topologies consistent with introgression across *Heliconius* chromosome 18 compared to the Z chromosome is only partially due to an elevated rate of gene flow on autosomes (Martin *et al*. 2013), but also reflects increased incomplete lineage sorting due to their larger effective population size relative to the sex chromosome. Weightings may be more difficult to interpret in cases with a less well understood evolutionary history. Nevertheless, it may be possible for example to differentiate between topologies representing the 'species tree' and those reflecting recent introgression by comparing their branch lengths (which can be output by *Twisst*) (Fontaine *et al*. 2015; Gante *et al*. 2016). Finally, we have found that topology weighting provides a means to identify candidate loci underlying trait variation, based on clustering of taxa by phenotype [see also Van Belleghem et al. (2017), for a more extensive demonstration of this power]. In summary, topology weighting is a simple but versatile exploratory tool that is applicable to a diverse range of questions and datasets.

## Acknowledgements

We thank Pasi Rastas, Markus Möst and Joe Hanly for useful comments on the method. SHM was funded by a research fellowship from St John’s College, University of Cambridge. SMVB was funded by NSF grant DEB-1257839. Additional funding was provided by European Research Council grant 339873 held by Chris Jiggins.

## Supplementary Figures

**Figure S1.**
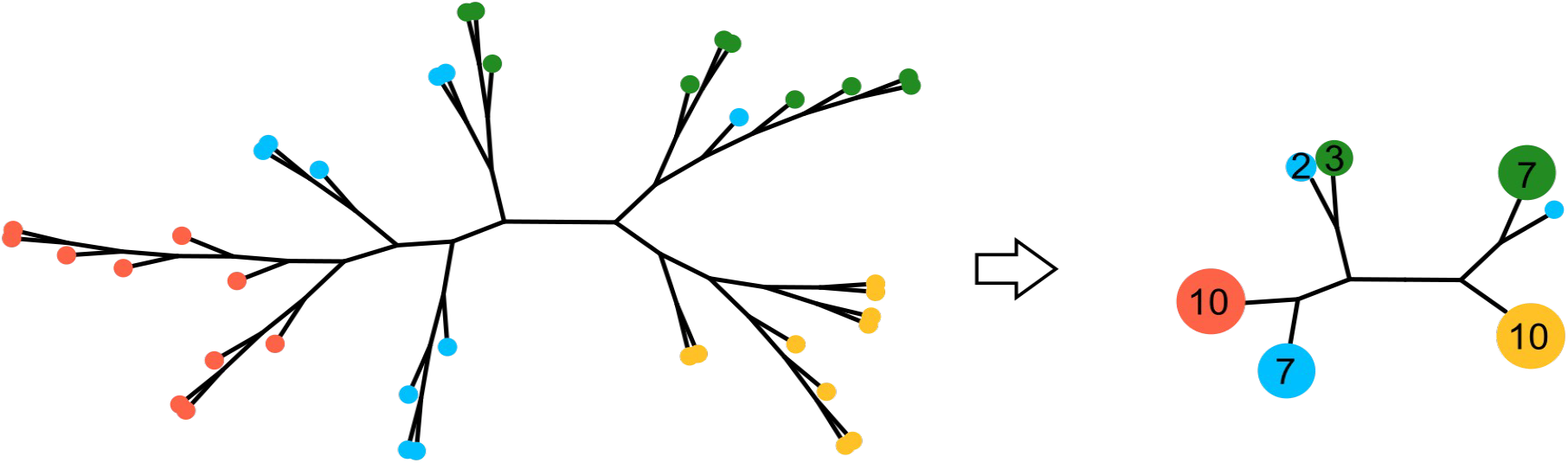
Tree Simplification. The full tree (left) has four defined taxa of ten samples each, indicated by different colours. This equates to 10,000 (10^4^) unique sub-trees that include a single individual from each taxon. *Twisst* reduces the number of unique sub-trees to count by collapsing clades and weighting them by the number of individuals of each taxon present (and adjusting branch lengths accordingly). In this example, after this process, there remain only 6 unique sub-trees to count (3x2x1x1).

**Figure S2.**
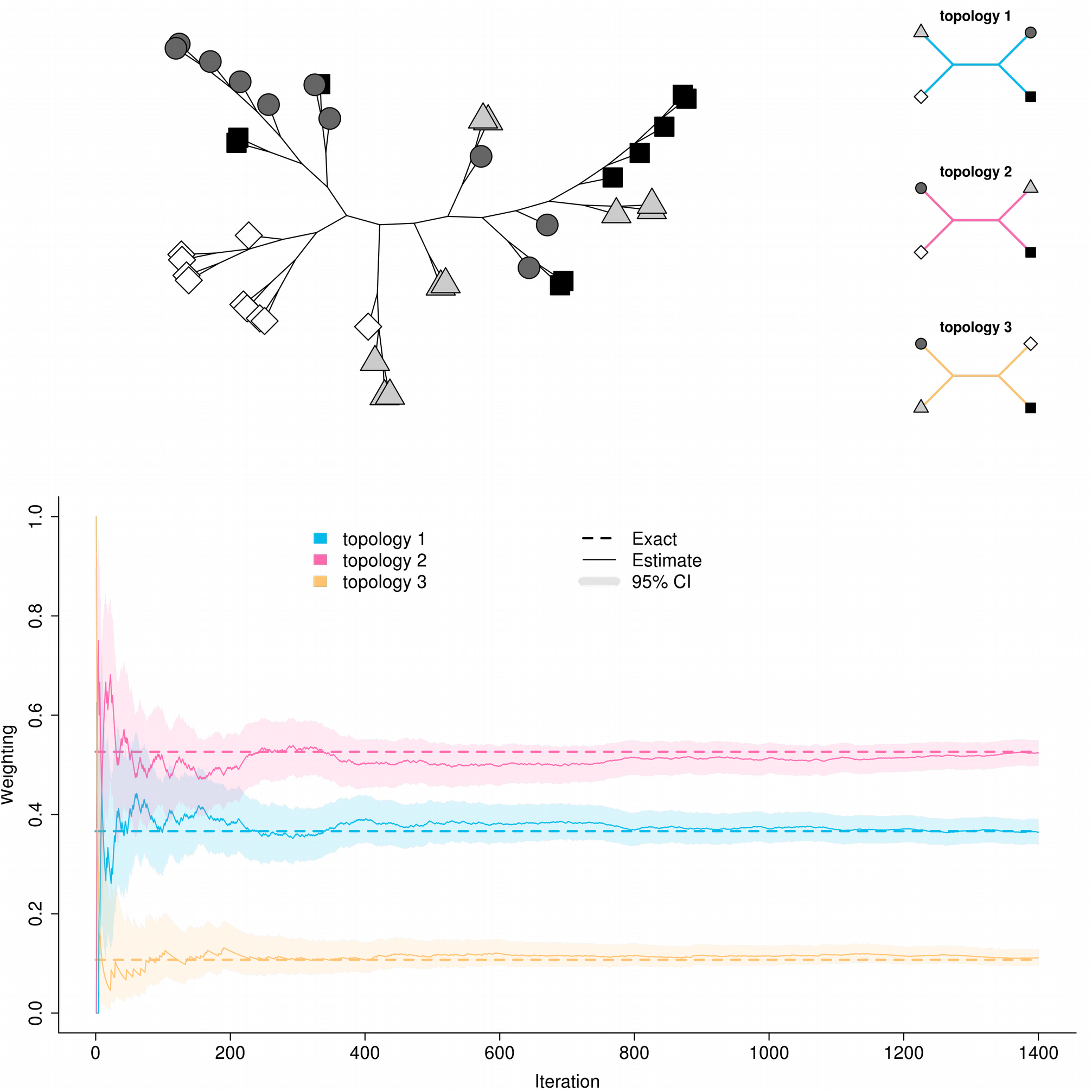
Approximate weighting by random sampling. This example tree has four defined taxa indicated by different symbols. There are three possible taxon topologies (top right). The graph shows the estimated weighting for each topology (Y-axis) after randomly sampling a number of sub-trees from the full tree (X-axis). The 95% binomial confidence interval (calculated using the Wilson method), is shaded. Dashed lines show the true exact weightings computed by sampling all 10,000 sub-trees.

**Figure S3.**
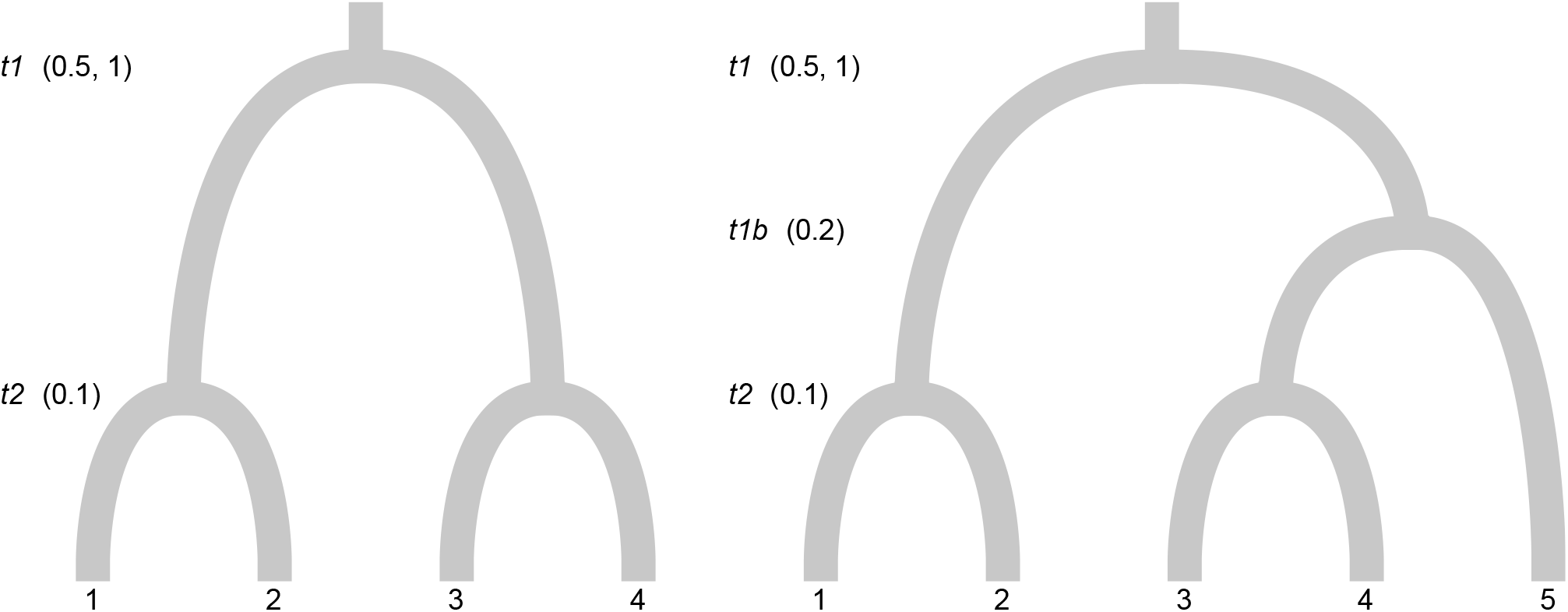
Demographic scenarios for power simulations. Four (left) and five (right) population simulations were performed. Split times are shown, note that two different values were tested for split time *t1* in both scenarios.

**Figure S4.**
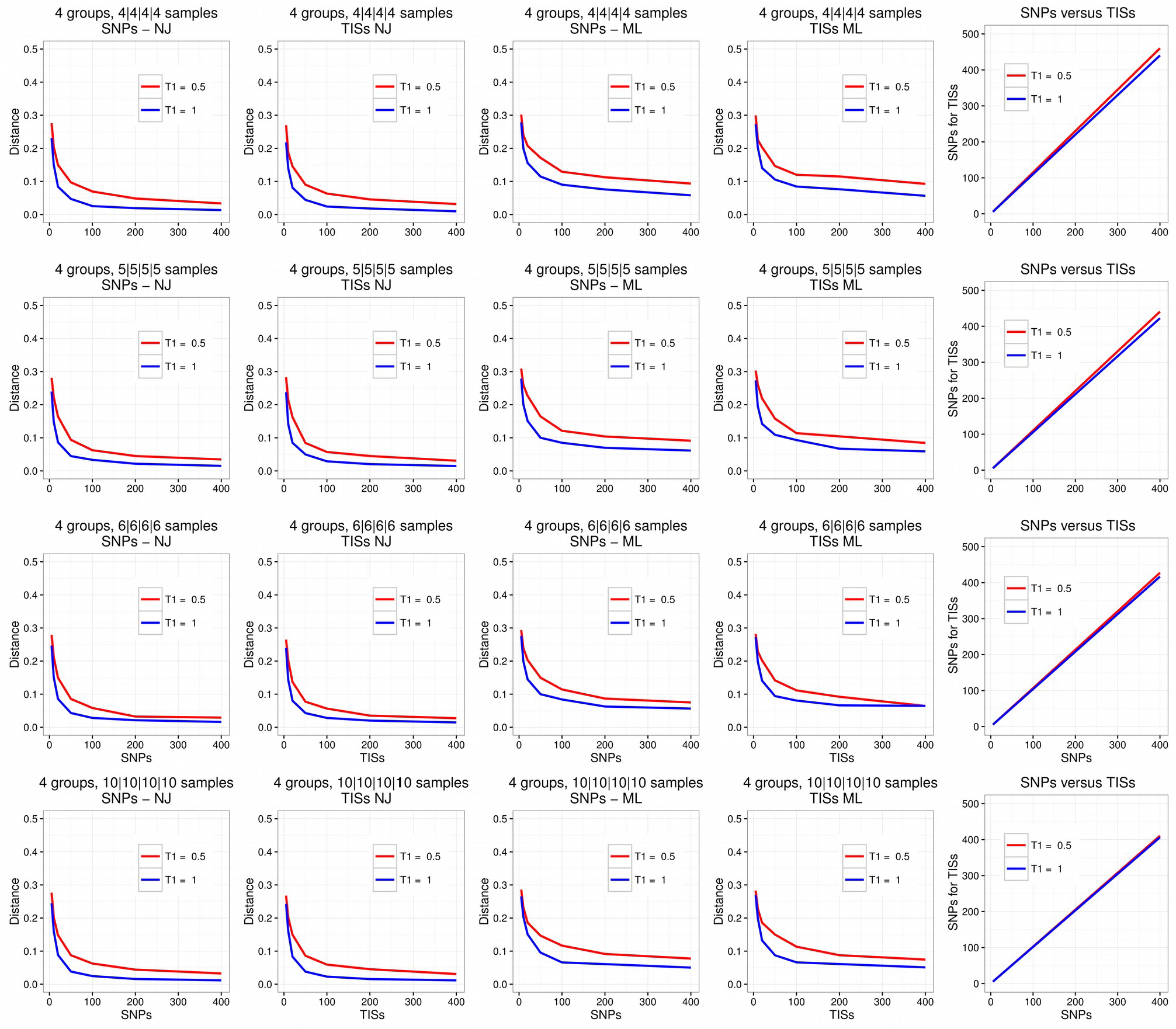
Power analyses with four equal-sized groups. Each plot in the first four columns shows the error rate, calculated as a scaled euclidean distance between the true and inferred weightings (Y-axis), plotted against the number of sites used for tree inference (X-axis). The simulated sequences were truncated to create a sequence of the correct length for tree inference. Truncation was performed either after X SNPs were observed (columns 1 and 3) or after X ‘taxon-informative sites’ (TISs) had been observed (columns 1 and 4). The final column gives the number of SNPs required (Y-axis) to observe a certain number of TISs (X-axis). The first two columns show results after tree inference using neighbour joining (NJ) and the next two columns show results after tree inference using maximum likelihood (ML). Each row represents a distinct sampling strategy (four groups of four samples, four groups of five samples etc.). Red and blue lines indicate the two different demographic scenarios tested, with a different split time *t1* (see Fig. S3).

**Figure S5.**
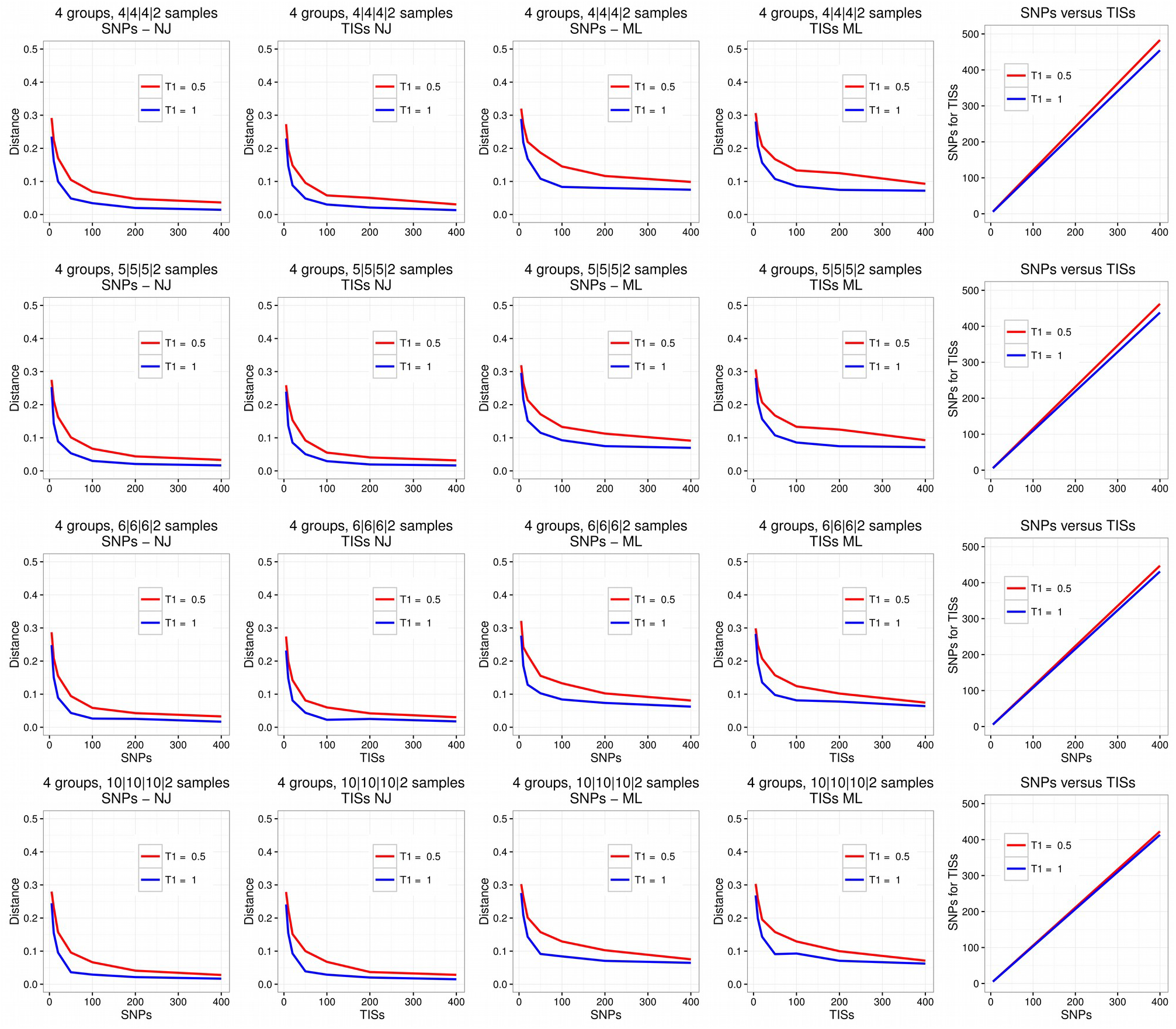
Power analyses with four groups of different sizes. As in Fig. S4, except for groups of different sizes.

**Figure S6.**
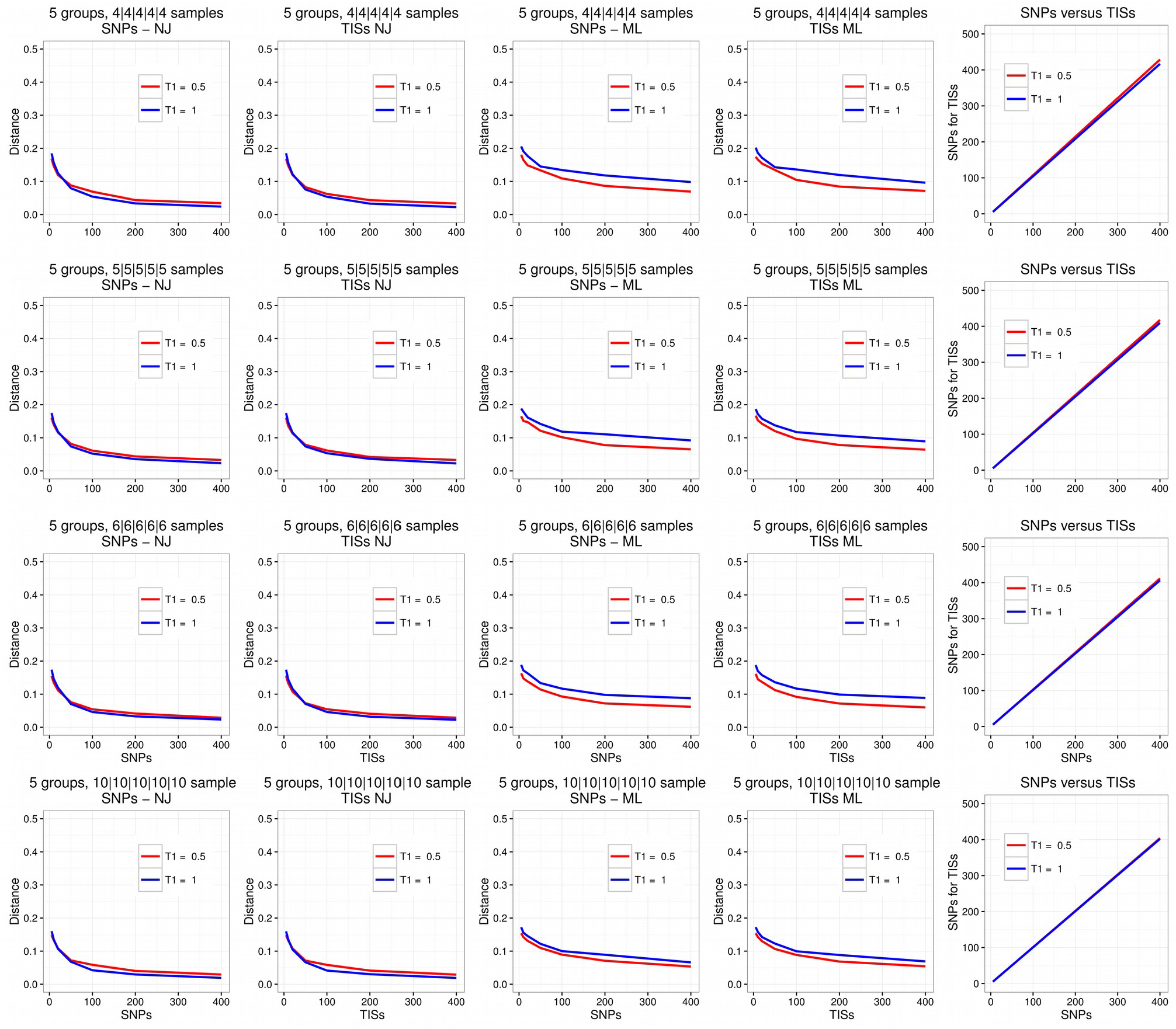
Power analyses with five equal-sized groups. As in Fig. S4, except for simulations with five groups.

**Figure S7.**
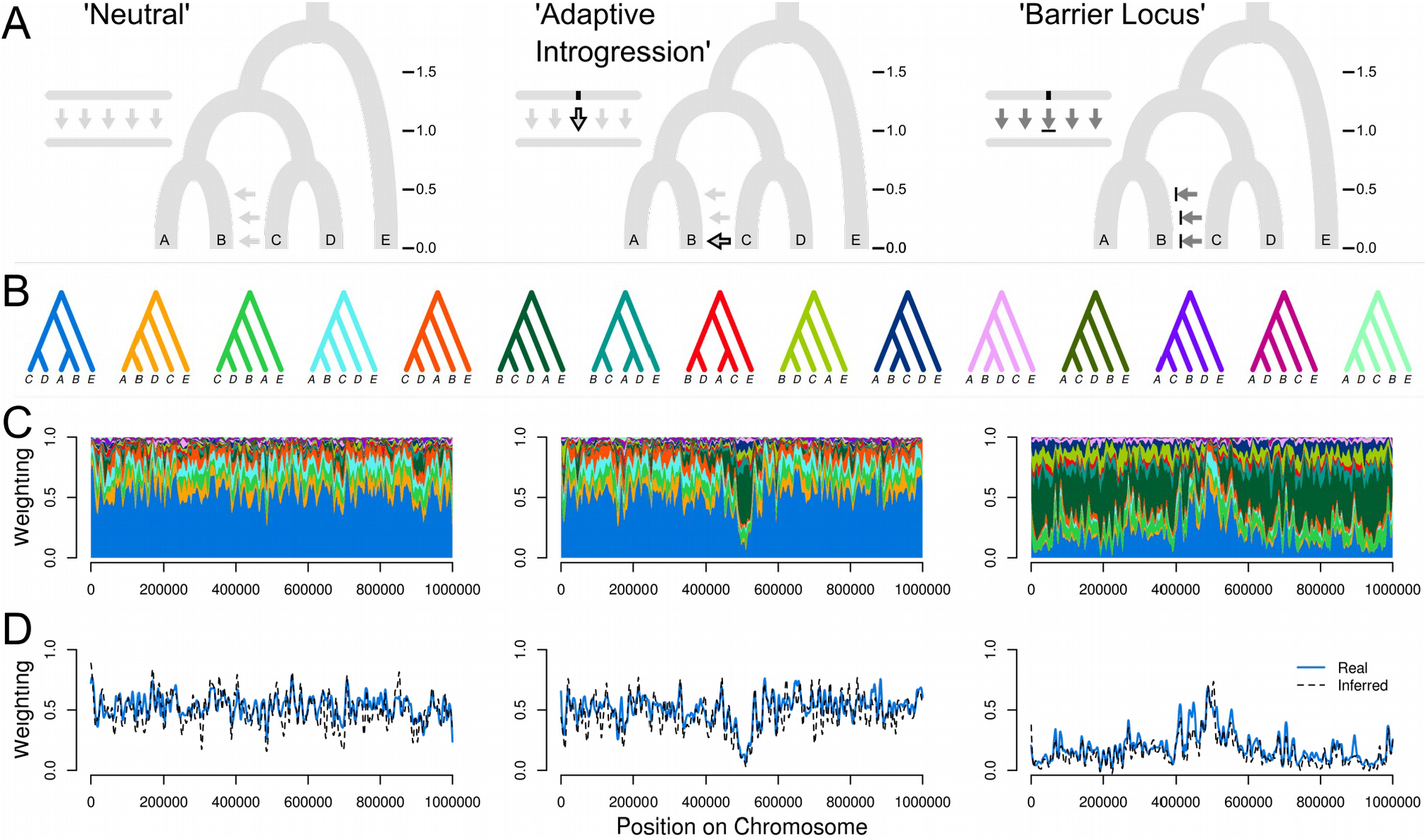
Tests on simulated chromosomes with 5 taxa. **(A)** In all three demographic scenarios, populations split in the order ((((A,B),C),D),E), at the split times indicated (in units of 4N generations), with migration from C to B. In the ‘Neutral’ scenario, there is no selection and moderate migration. The ‘Adaptive Introgression’ scenario is similar, except a beneficial allele at a locus in the centre of the chromosome is allowed to move from population C into B at time 0.1. In the ‘Barrier Locus’ scenario, the rate of migration is high, but an allele at the central locus that is fixed in C is selected against in population B. **(B)** All fifteen possible taxon topologies. Note that we illustrate the topologies for the four taxa as rooted, with E as the outgroup for simplicity, but the rooting is not considered when computing the weightings. **(C)** Weightings for all topologies plotted (stacked) across the chromosome, with loess smoothing (span = 20 kb). (D) Weightings for topology (((A,B),C),D) inferred from simulated sequence data using non-overlapping 50 SNP windows and neighbour joining. Solid blue lines indicate the true values, and dashed black lines indicate the inferred values. Values are smoothed as above.

**Figure S8.**
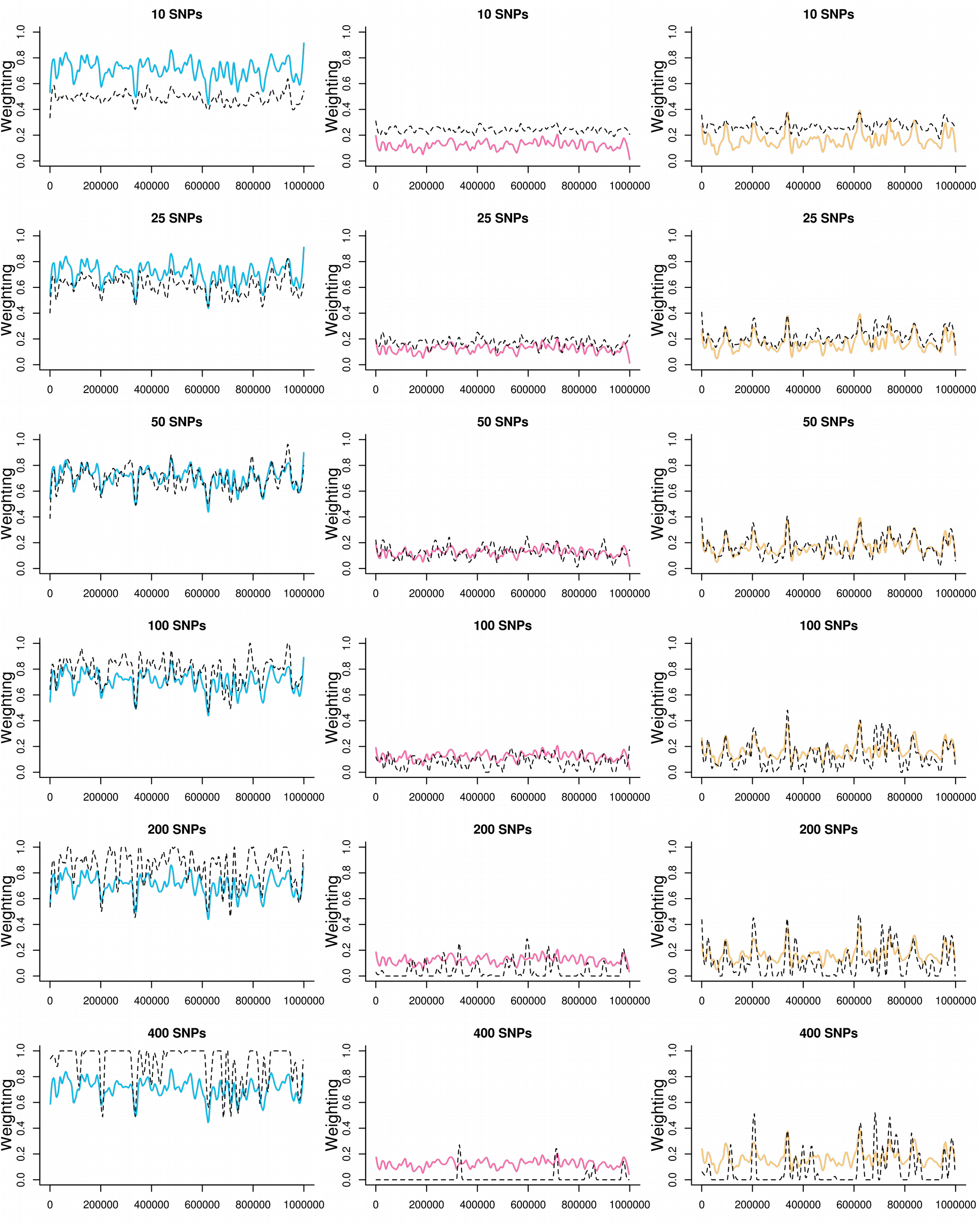
Inferred vs true weightings using different window sizes (ρ = 0.01) The true (solid coloured line) and inferred (dashed black line) weightings, plotted across the simulated 1 Mb chromosome, with loess smoothing (span 0.04). The three columns with different colours represent the three taxon topologies (see Fig. 2 in the main paper). Rows represent different window sizes (fixed number of SNPs) used for tree inference (neighbour joining).

**Figure S9.**
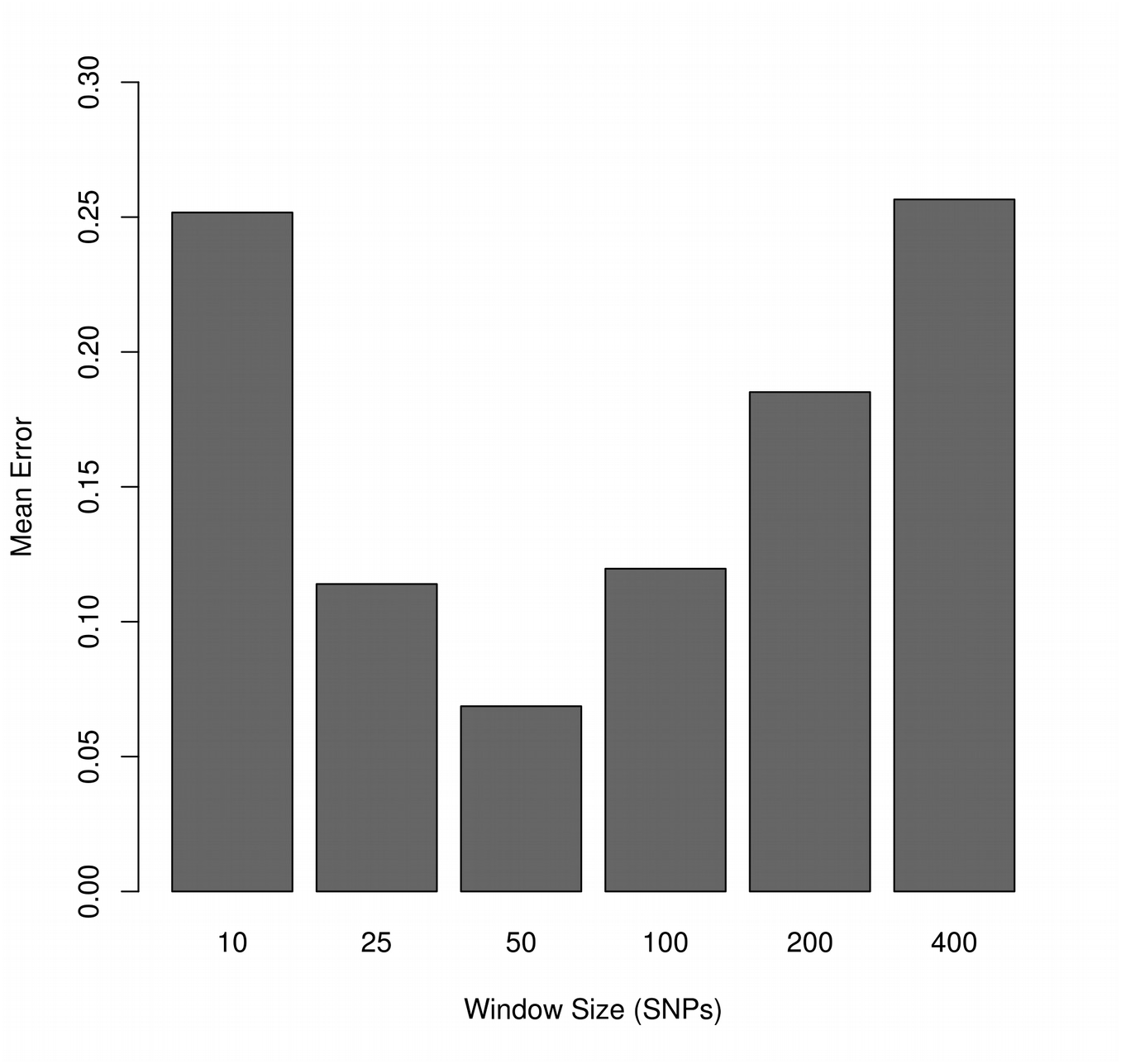
Weighting error rates using different window sizes (ρ = 0.01) Error rate, calculated as a scaled euclidean distance between the true and observed weightings, averaged over the 1 Mb simulated shromosome. Error rates were computed after first smoothing both the observed and true weightings using loess (span = 0.04).

**Figure S10.**
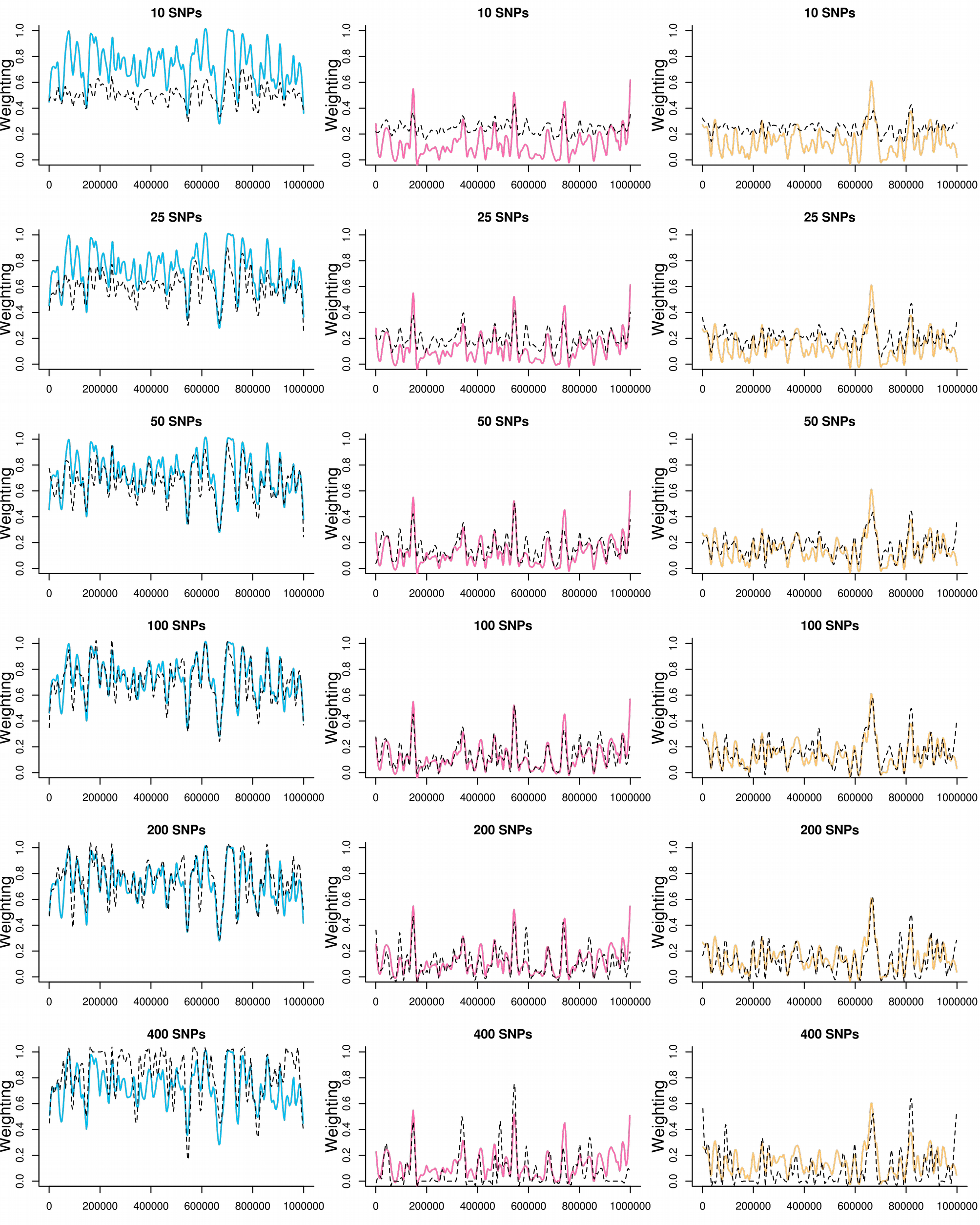
Inferred vs true weightings across simulated chromosomes (ρ = 0.001) As in Fig. S8, except here simulations used a population recombination rate (ρ) of 0.001.

**Figure S11.**
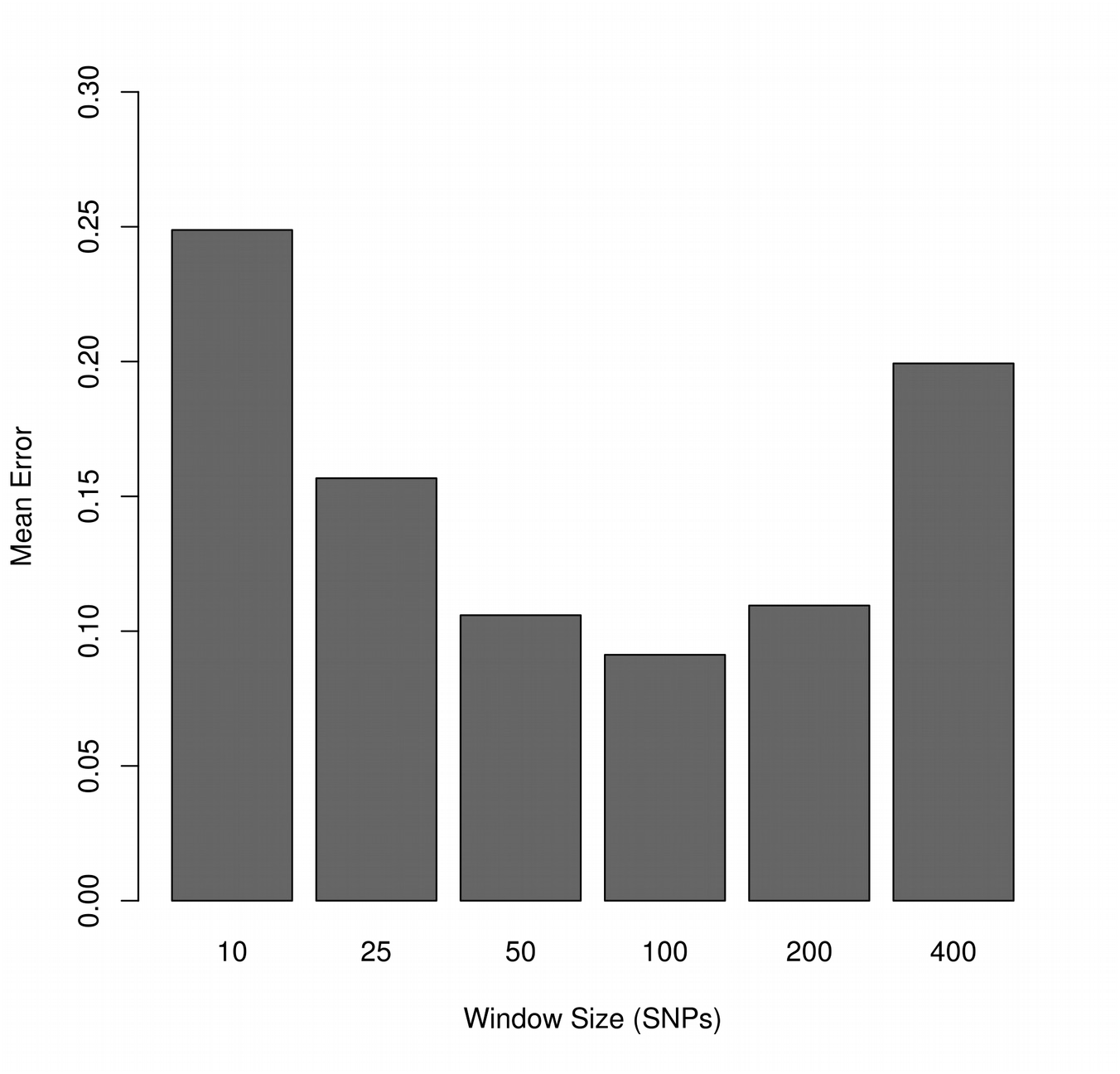
Weighting error rates using different window sizes (ρ = 0.001) As in Fig. S9, except here simulations used a population recombination rate (ρ) of 0.001.

**Figure S12.**
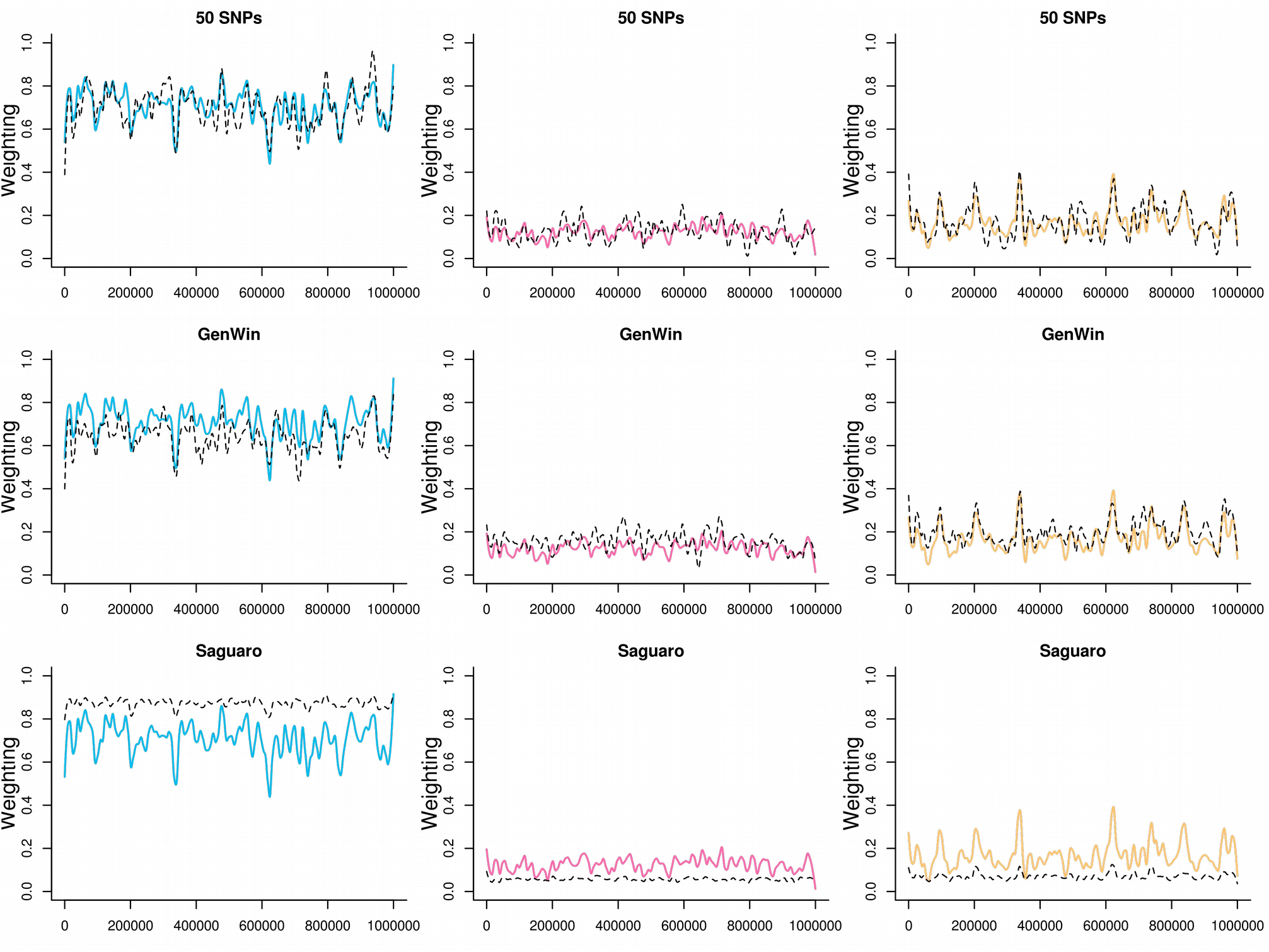
Inferred vs true weightings across simulated chromosomes. As in Fig. S8 and S10, except here comparing different methods for window-based tree inference: 50 SNP windows (first row), *WinGen* inference of breakpoints (second row) or *Saguaro* (row three).

**Figure S13.**
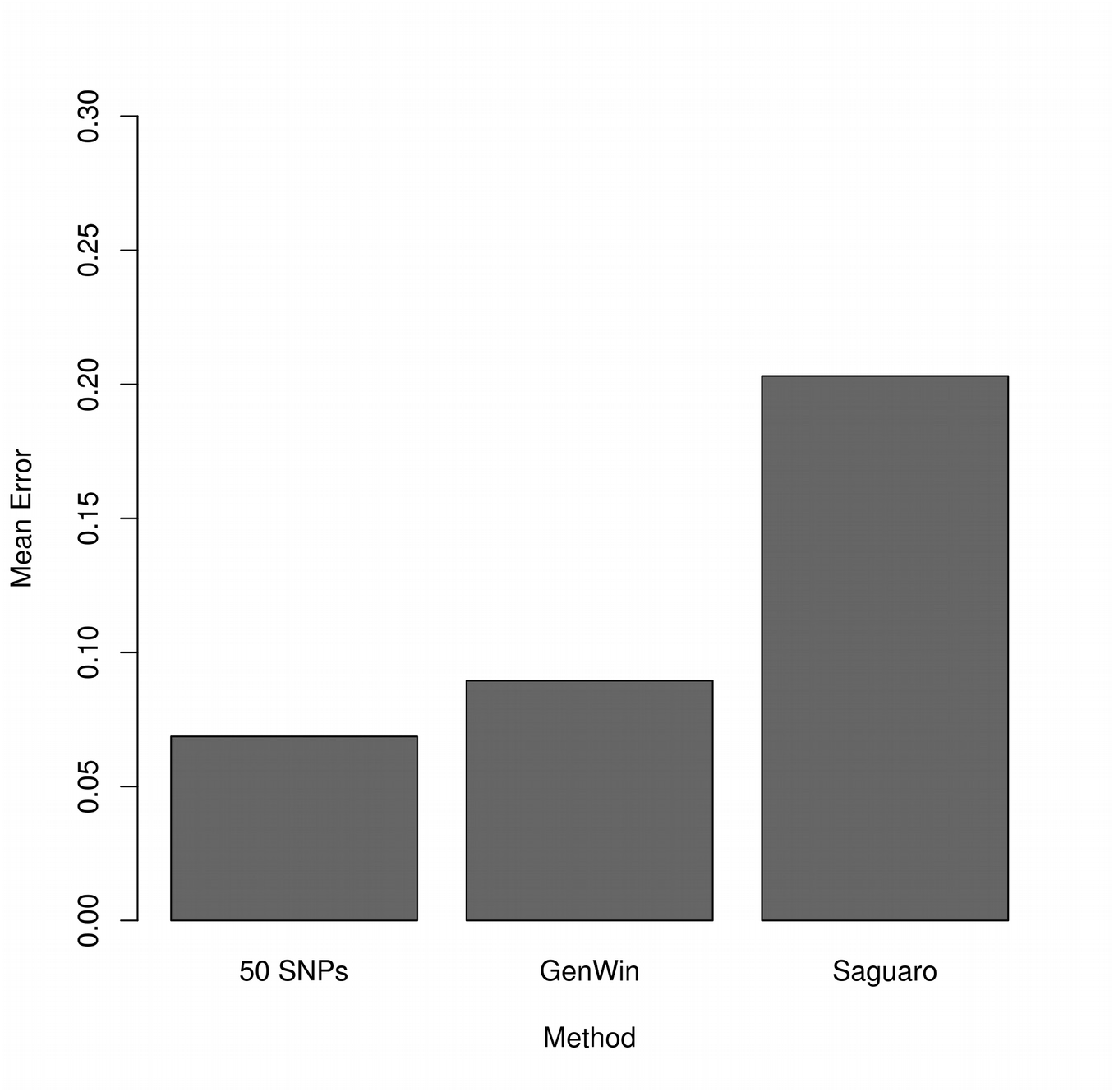
Weighting error rates using different methods for window-based tree inference. As in Fig. S9 and S11, except here comparing different methods for window-based tree inference.

**Figure S14.**
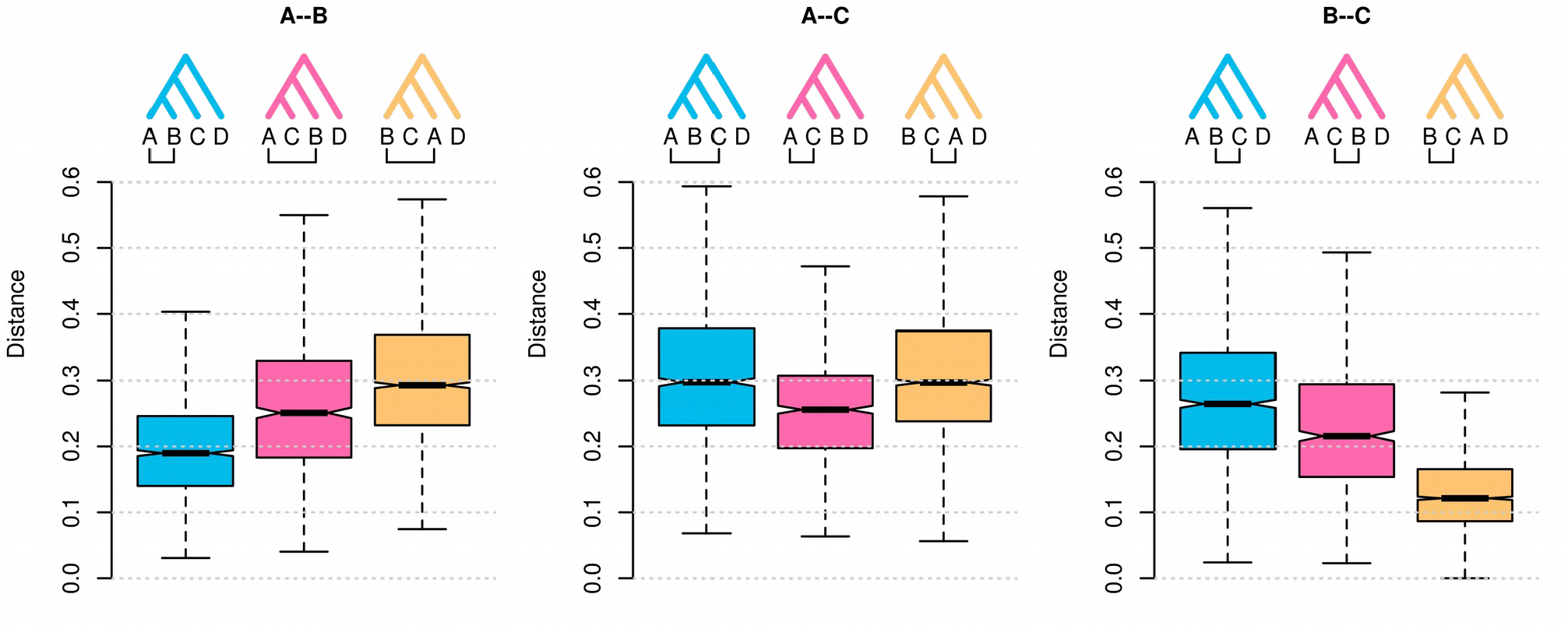
Average branch length separating each pair of taxa in different sub-trees. Boxplots show the distribution of average pairwise distances (i.e. the branch lengths separating each pair of taxa) across all trees, separated by the topology matched by each sub-tree (shown above and colored). For example, the blue box in the left-hand plot gives the distribution of average pair-wise distances between samples from taxon A and B for all sub-trees that matched the topology (((A,B),C),D). The corresponding distances are plotted for the other two topologies in different colors, and then for the other pairwise comparisons (A-C and B-C) in the middle and right-hand plots, respectively. The lower average distance between sequences from B and C (right-hand plot) in topologies where they coalesce first (yellow), compared to the distance between B and A (left-hand plot) in topologies where they coalesce first (blue), suggests that the yellow topology results from recent introgression between B and C, whereas the blue topology matches the population branching order.

**Figure S15.**
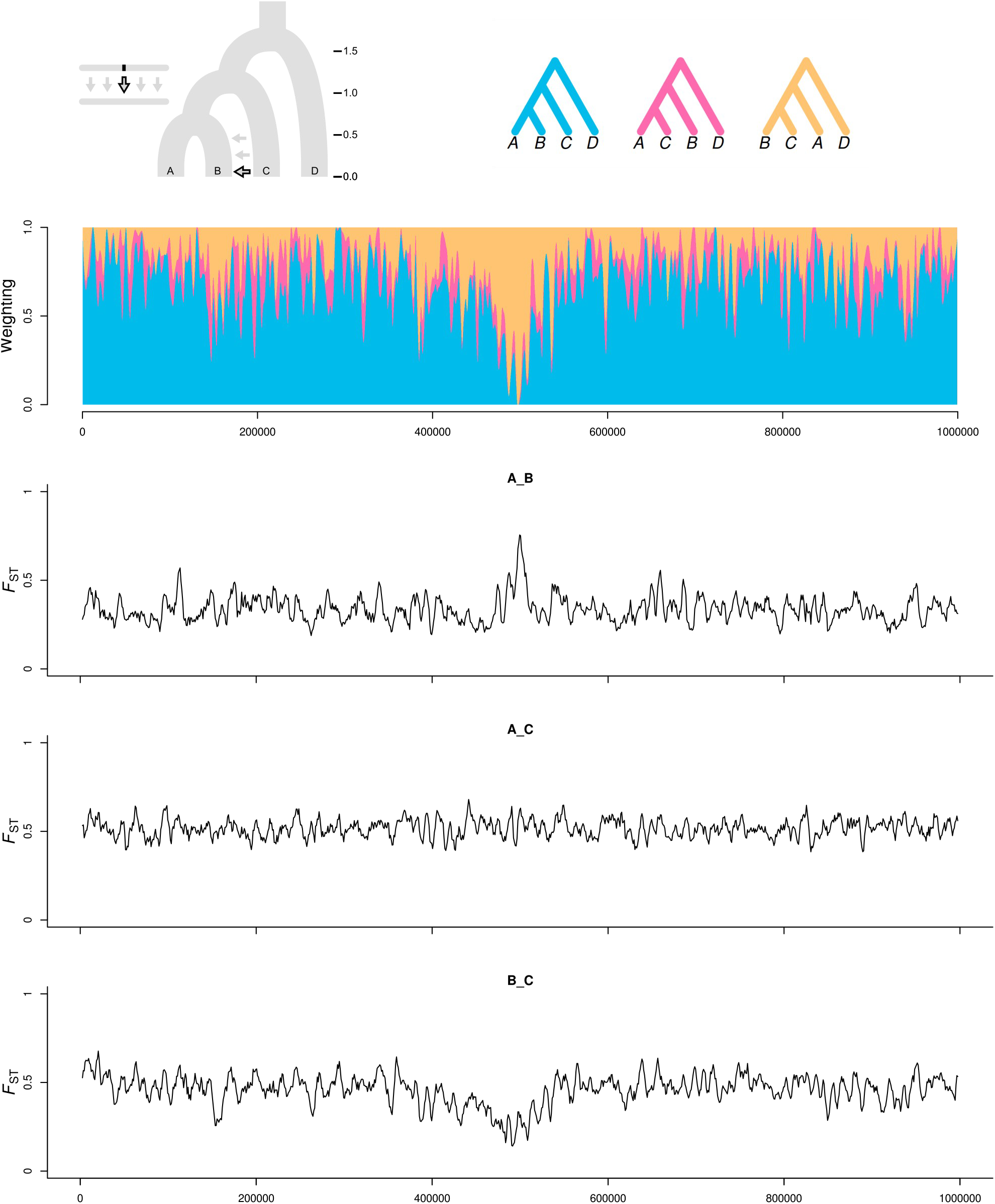
Comparison between topology weighting and *F*_ST_. The simulated ‘Adaptive Introgression’ (top) scenario was used to compare topology weighting to *F*_ST_. Weightings are plotted as in Fig. 2B except with a loess smoothing (span = 10 kb). Pairwise *F*_ST_ plots for each pair of ingroup taxa are plotted for 5 kb sliding windows, moving in increments of 1 kb. *F*_ST_ captures the signal of the adaptive introgression between populations B and C in the form of a peak of divergence between A and B, and marginally reduced divergence between B and C.

**Figure S16.**
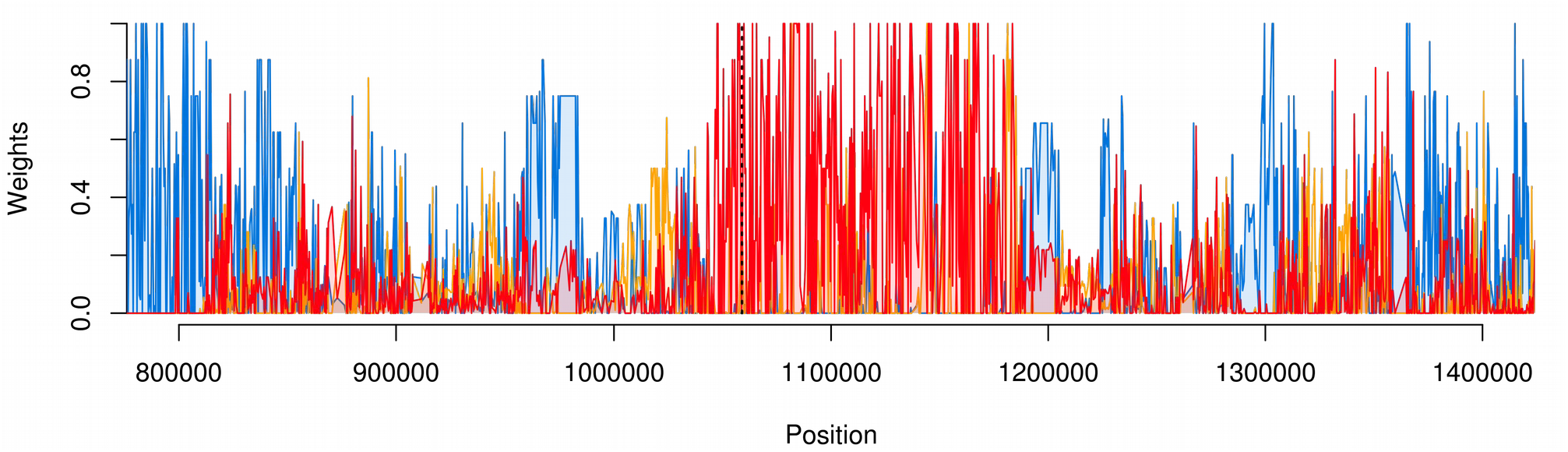
Topology weighting around the *Heliconius optix* gene. Unsmoothed weightings for three topologies from the *Heliconius* analysis, topo3 (blue), topo6 (orange) and topo11 (red) (see Fig. 4 in the main paper for details), are plotted across the region of Chromosome 18 around the gene *optix* (indicated by a dashed black line). A ∼150 kb block of high weightings for topo11, which supports introgression between *H. melpomene amaryllis* and *H. timareta thelxinoe*, includes *optix* and its downstream regulatory region that is known to control red wing patterning.

